# TCR-pMHC complex formation triggers CD3 dynamics

**DOI:** 10.1101/2022.07.27.501668

**Authors:** Floris J. van Eerden, Alrahman Aalaa Sherif, Mara Anais Llamas-Covarrubias, Arthur Millius, Xiuyuan Lu, Shigenari Ishizuka, Sho Yamasaki, Daron M. Standley

**Affiliations:** Department of Systems Immunology, Immunology Frontier Research Center, Osaka University, 3-1 Yamadaoka, Suita, 565-0871, Japan; Department of Genome Informatics, Research Institute for Microbial Diseases, Osaka University, 3-1 Yamadaoka, Suita, 565-0871, Japan; Laboratory of Molecular Immunology, Immunology Frontier Research Center, Osaka University, 3-1 Yamadaoka, Suita, 565-0871, Japan; Department of Molecular Immunology, Research Institute for Microbial Diseases, Osaka University, 3-1 Yamadaoka, Suita, 565-0871, Japan

**Author notes:** Equal contribution E-mail: Floris van Eerden Daron Standley.

## Abstract

In this study, we present an allosteric mechanism for T cell receptor (TCR) triggering upon binding a peptide-MHC complex (pMHC), in which a conformational change in the TCR upon pMHC binding controls the mobility of the CD3 proteins. We found that the TCRβ FG loop serves as a gatekeeper, preventing accidental triggering, while the connecting peptide acts as a hinge for essential conformational changes in the TCR. Atomistic simulations and cell-based experiments with genetically modified connecting peptides demonstrate that rigidified hinge residues result in excessive CD3 dynamics and hypersensitivity to pMHC binding. Our model thus provides a clear connection between extracellular TCR-pMHC binding and changes in CD3 dynamic that propagate from outside to inside the cell.

## Introduction

T cell receptors (TCRs) specifically recognize peptide antigens presented by major histocompatibility complex (MHC) molecules. The TCR is a heterodimer composed of an ɑ and a β chain that forms a complex with three CD3 dimers (CD3γε, CD3δε and CD3ζζ) (Figure 1A). The binding of agonistic, but not antagonistic, peptide-MHC (pMHC) complexes results in Lck-mediated phosphorylation of immunoreceptor tyrosine-based activation motifs (ITAMs) ^1^. The ITAMs, whose phosphorylation is required for T cell activation, are located on the cytoplasmic tails of the CD3s. Current evidence suggests that the CD3 ITAMs are hidden in the membrane in the resting state, but are exposed to the cytoplasmic kinase Lck upon pMHC binding^2, 3^. The effector functions of activated T cells range from killing infected cells to inhibition of immune responses to establishment of memory of specific peptide antigens. Although TCRs have been intensively studied for several decades, the mechanisms linking pMHC engagement to TCR triggering remain elusive. It has been difficult to rationalize how TCR-pMHC binding could result in changes in the cytoplasmic regions, since the TCR chains lack a proper cytoplasmic domain, while the CD3 proteins are too short to interact with the pMHC. The fact that the TCR and CD3 associate non-covalently further complicates the problem.

**Figure 1:**
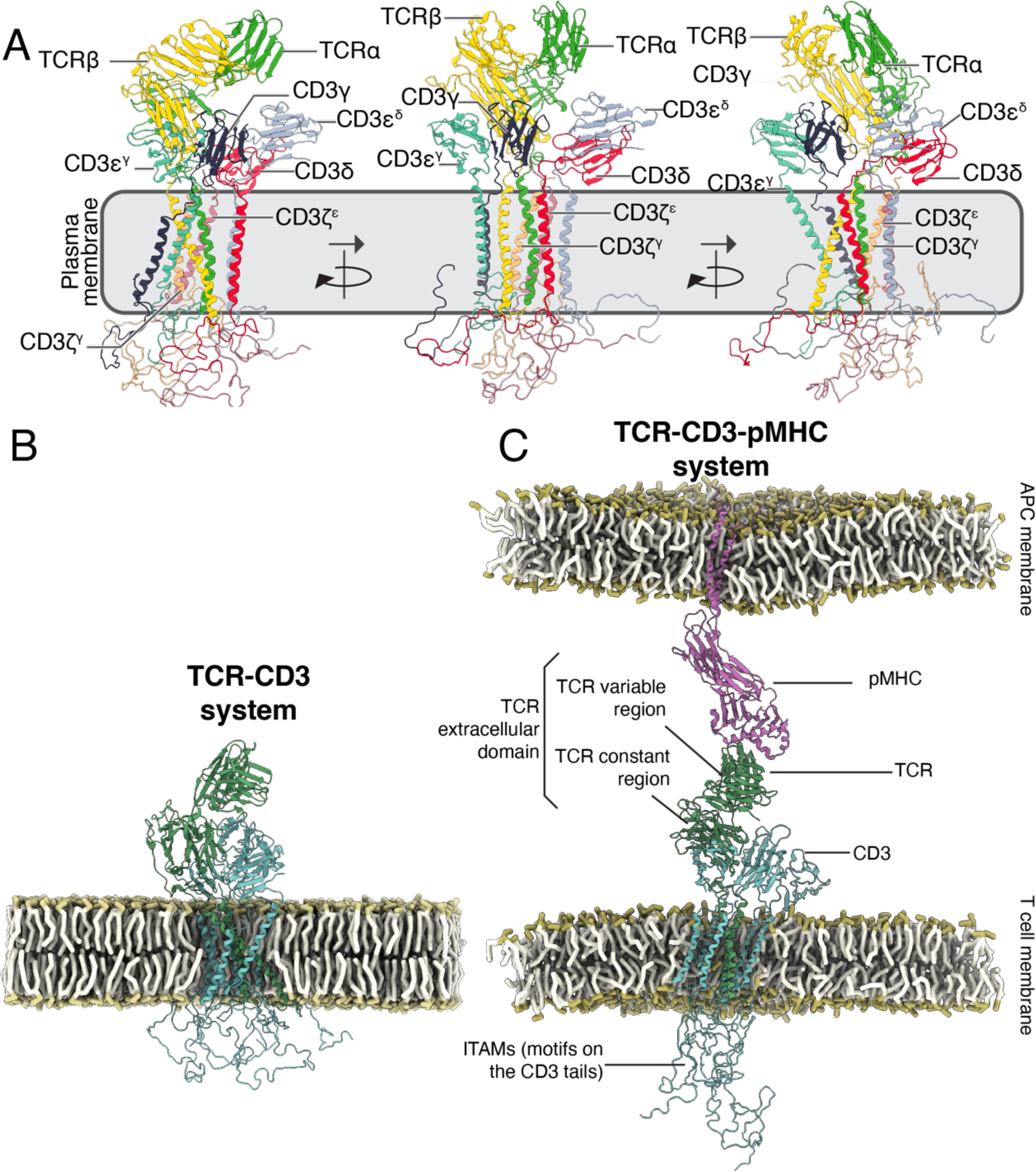
The TCR forms a complex with CD3 proteins. Overview (A) of the TCR and the CD3 proteins. A superscript is used to distinguish the CD3ε and CD3ζ chains. CD3ε^δ^ refers to the CD3ε from the CD3δε dimer and likewise CD3ε^γ^ refers to the CD3ε of the CD3γε dimer. CD3ζ^γ^ refers to the CD3ζ closest to the transmembrane (TM) region of CD3γ and CD3ζ^ε^ to the CD3ζ closest to the TM region of CD3ε^δ^. Structure from PDB code 6JXR^15^. Snapshots of the TCR-CD3 (B) and TCR-CD3-pMHC (C) simulation systems. The TCR (green), CD3 proteins (cyan), pMHC (purple), and lipid headgroups (light brown) are shown. In the TCR-CD3-pMHC system a pore is present in the upper membrane to allow the flow of solvent between the two compartments, which is why headgroup beads are visible in the membrane interior. Solvent and lipid tails are not shown for clarity. In (C), the TCR variable and constant regions are indicated, as well as the ITAMs on the CD3 proteins. The TCR extracellular domain (EC) encompasses the variable and constant regions.

In the last decade, a number of models of TCR triggering and T cell activation, each emphasizing different aspects of the phenomenon, have been proposed. The relationship between initial pMHC engagement and cytoplasmic signaling is described by the *mechanosensor* model, in which TCRs discriminate agonistic from antagonistic pMHCs via catch bond formation^4–7^. In this model, pMHC binding transfers force from the pMHC-TCR interface to the TCR-CD3 interface via the TCRβ-FG loop^8, 9^. In the *kinetic proofreading* model, the binding lifetime of the pMHC-TCR complex enables differentiation of agonistic pMHCs from antagonistic ones^10^. The *TCR bending mechanosignal* model argues that TCR-pMHC engagement alters membrane curvature, which, in turn, promotes conformational changes in the receptors that result in TCR triggering^11^. Alternatively, in the *allosteric relaxation* model, TCR-pMHC binding decouples TCR and CD3ζ interactions^12–14^. Importantly, each of these models requires a yet unknown coupling between the TCR and CD3 that must ultimately be transmitted to the cytoplasm. The coupling and transmission are crucial because the CD3 chains – not TCRɑβ – contain the ITAMs.

In 2019, the first full-length TCR-CD3 complex structure was determined by single particle cryo-EM^15^. Interestingly, the extracellular domain (EC) of the TCR was bent with respect to the membrane. Subsequent molecular dynamics simulations of the TCR showed that the angle between the TCR EC and transmembrane domain (TM) is dynamic, allowing the TCR EC to adopt a range of different conformations^16, 17^. In 2022, the structure of a pMHC-bound TCR-CD3 complex revealed that the TCR was unchanged by the binding to a soluble pMHC ^18^. We reasoned that, since the static structures of pMHC-bound and unbound TCRs do not reveal an obvious triggering mechanism, the dynamics of the complex might hold clues. To this end, we used extensive multi-scale molecular dynamics simulations to investigate the interplay of TCR, CD3 and pMHC. The resulting picture is one in which CD3 dynamics plays a central role in distinguishing between bound and unbound TCRs.

### Description of the simulation systems

To study the dynamics of the TCR under resting and pMHC-engaged conditions, two simulation systems were constructed: a TCR-CD3 system and a TCR-CD3-pMHC system. The TCR-CD3 system consisted of a single TCR-CD3 complex embedded in a POPC bilayer (Figure 1B), while the TCR-CD3-pMHC system was composed of two opposing POPC bilayer slabs with the lower containing a TCR-CD3 complex and the upper containing a pMHC (Figure 1C). The simulations were performed before the recently reported TCR-CD3-pMHC structure was published^18^; therefore, to model the complex, we created a chimeric TCR in which the TCR transmembrane (TM) region from the De Dong (PDB: 6jxr)^15^ structure was docked onto the pMHC-TCR complex structure by Newell (PDB: 3qiu)^19^. Harmonic potentials centered on the crystal structure were used to tether the TCR to the pMHC; the potentials were applied in a stepwise manner during 1.5 µs of preparatory simulations to allow the system to incrementally adjust (Figure S1). A pore was created in the pMHC-bound membrane to allow the flow of solvent between the two membrane compartments; this flow was required to permit the distance between the two membranes to fluctuate naturally. For each system, one hundred 10 µs production runs were performed, totaling 1 ms per system.

*CD3ε chains manifest a wider range of motion in the TCR-CD3-pMHC than the TCR-CD3 system* Iso-occupancy surfaces were calculated to examine the distribution of the CD3ε chains in the two systems. In the TCR-CD3-pMHC simulations, the CD3ε chains sampled a larger area around the TCR compared to the TCR-CD3 simulations (Figure 2). In the TCR-CD3 simulations, the iso-occupancy surface of the CD3ε chains had a clear minimum close to the position of the CD3ζ chains (Figure 2A). By contrast, in the TCR-CD3-pMHC simulations, the CD3ε chains completely sampled the region around the TCR (Figure 2B). Interestingly, the iso-occupancy surface in the TCR-CD3 simulation was not symmetric – the iso-occupancy was particularly low around the CD3ζ chain adjacent to CD3γ (CD3ζ^γ^). This difference was also apparent when the CD3ε chain position was expressed in polar coordinates and the density plotted in a plane parallel to the membrane: The TCR-CD3 simulations indicated an excluded region from 225-315°, which was occupied in the TCR-CD3-pMHC simulations. Taken together, these results indicate that TCR-pMHC binding results in a greater range of CD3ε motion.

**Figure 2:**
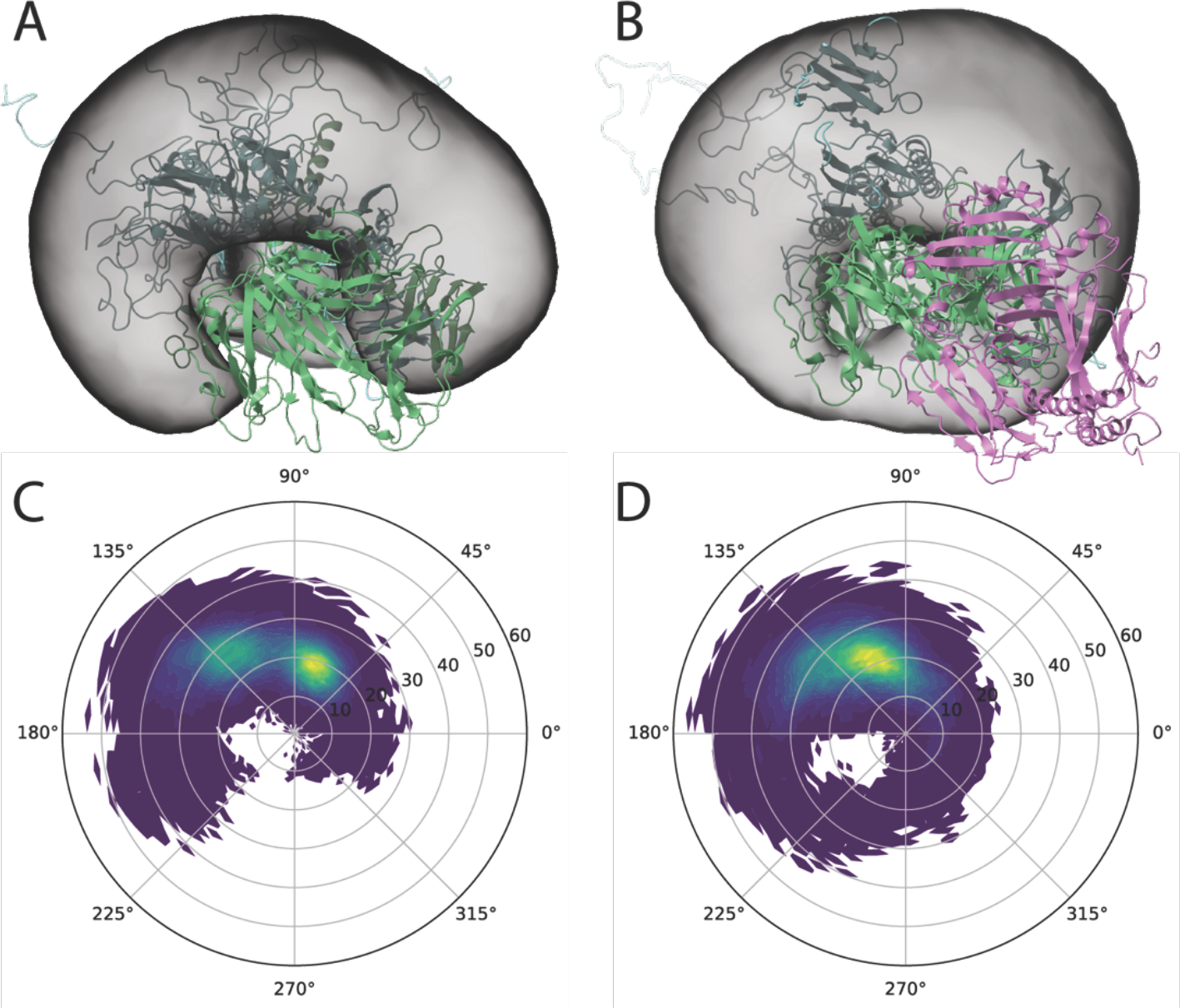
Distribution of CD3ε chains around TCR αβ visualized by iso-occupancies and polar plots. Iso-occupancy of CD3ε^γ^ε^δ^ in the TCR-CD3 simulation (A) and in the TCR-CD3-pMHC simulation (B). The TCR (lime), CD3 (cyan), which are mostly obscured by the iso-surfaces, and pMHC (purple) are shown. Polar plots of CD3ε^γ^ chains around TCRβ chain in the TCR-CD3 simulation (C) and in the TCR-CD3-pMHC simulation (D).

*TCR-pMHC engagement resulted in fewer contacts between CD3 and the TCRβ variable domain* To understand the difference in the CD3 spatial distributions in the two simulations, we next analyzed contacts between the CD3 and TCRɑ/β chains. In the TCR-CD3 simulations, the TCRβ variable domain was constantly in contact with the CD3γε dimer, but these interactions virtually disappeared in the TCR-CD3-pMHC simulations. In the TCR-CD3 systems, the TCRβ variable domain made on average 7445 (stdev 7215) contacts per simulation with CD3ε^γ^ – the CD3ε next to CD3γ. In addition, the TCRβ variable domain made on average 1581 (stdev 2848) contacts per simulation with CD3γ. In the TCR-CD3-pMHC simulations, the number of contacts dropped to 195 (stdev 625) and 3 (stdev 14) for CD3ε^γ^–TCRβ and CD3γ–TCRβ variable domain contacts, respectively (Figure 3). These observations are consistent with a model in which TCR-CD3 contacts restrict CD3 movement.

**Figure 3:**
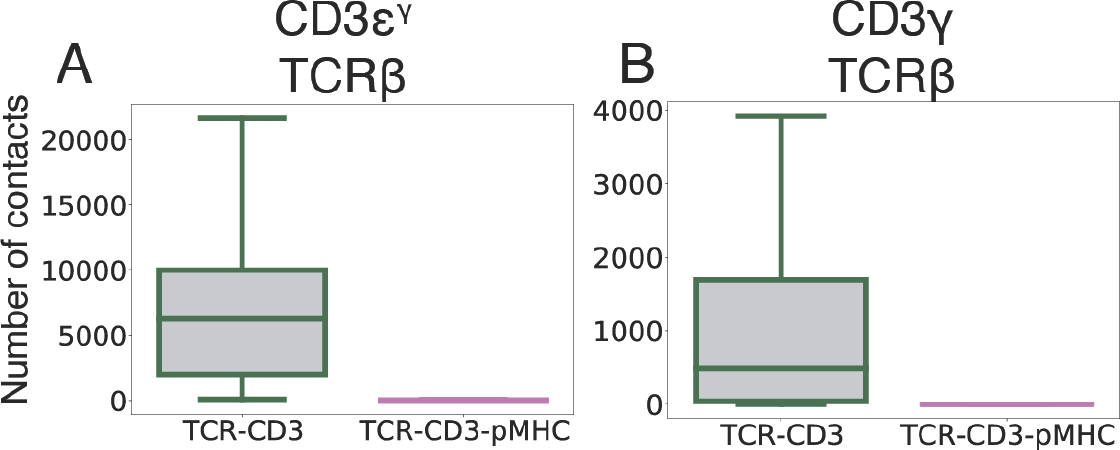
TCR-CD3 contacts. The number of contacts between the TCRβ variable domain and the indicated CD3 chains in the TCR-CD3 (green) and TCR-CD3-pMHC (purple) simulations. For clarity outliers are not shown. Contacts per residue are shown in Figure S2.

### The TCR is more extended in the TCR-CD3-pMHC complex

To understand the structural basis for the loss of CD3-TCR contacts and increase in CD3 dynamics upon pMHC binding, we next examined the conformation of the TCR in the two simulations. The TCR adopted a different orientation relative to the membrane in the TCR-CD3 and TCR-CD3-pMHC systems, as measured by the TCR EC tilt, the angle between the TCRβ transmembrane domain (TM) and the extracellular domain (EC) (Figure 4A). In the TCR-CD3 system, the TCR EC folded over the CD3 proteins, and the TCR adopted predominantly bent conformations (smaller tilt angles) with a peak in the angular distribution of approximately 104° (Figure 4B). Such TCR bending has been observed in previous TCR-CD3 simulations^17^. In contrast, in the TCR-CD3-pMHC system, the TCRs had more extended and open conformations (larger tilt angles) with a peak in the angular distribution of approximately 150°, which was close to the initial angle observed in the cryo-EM structures. Thus, upon pMHC binding, the TCR EC tilted away from the CD3 proteins (Figure 4C). Furthermore, the width of angular distributions was narrower in the TCR-CD3-pMHC simulations with a full-width at half-maximum (FWHM) of approximately 30°, compared to that in the TCR-CD3 simulations (FWHM 40°). These observations were consistent with the decrease in contacts and increase in dynamics in the TCR-CD3-pMHC systems compared with the TCR-CD3 simulations.

**Figure 4:**
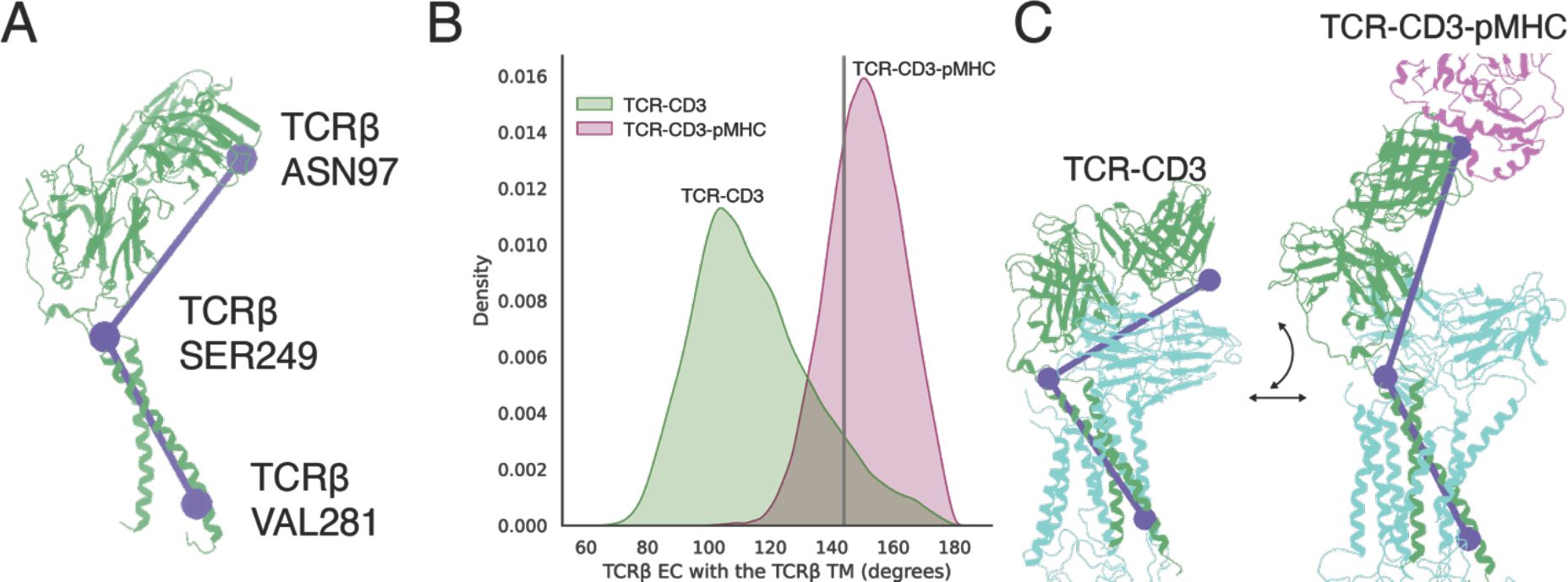
The TCR adopts a different conformation in the TCR-CD3 and TCR-CD3-pMHC simulations. (A) Illustration of the atoms used for the tilt angle calculation, which were TCRβ asparagine 97 TCRβ serine 249 and TCRβ valine 281. (B) Distribution of the tilt of the TCR EC (TCRβ EC-TCRβ TM angle) in the TCR-CD3 (green) and TCR-CD3-pMHC (purple) systems. The TCR EC tilt in the published cryo-EM structure^15^ (vertical line) was 144°. (C) Snapshots of the TCR-CD3 (left) and TCR-CD3-pMHC (right) simulations. Note the difference in the tilt angle. The TCR (green), CD3 proteins (cyan) and part of the pMHC (purple) are shown.

### The TCR-pMHC complex acts as drawbridge for the CD3 proteins

These observations suggest that the binding status of the TCR affects its conformation, which, in turn, affects CD3ε movement. Upon pMHC binding, the TCR adopts a more extended conformation, which breaks contacts between the TCR variable region and the CD3 proteins, allowing the CD3ε chains to diffuse around the TCR. In this manner, the TCR-pMHC complex acts as a “drawbridge” that releases the CD3 proteins, resulting in their increased movement (Figure 5). It is reasonable to hypothesize that, over time, the increase dynamics upon drawbridge formation is transmitted to the ITAMs, ultimately resulting in their phosphorylation.

**Figure 5.**
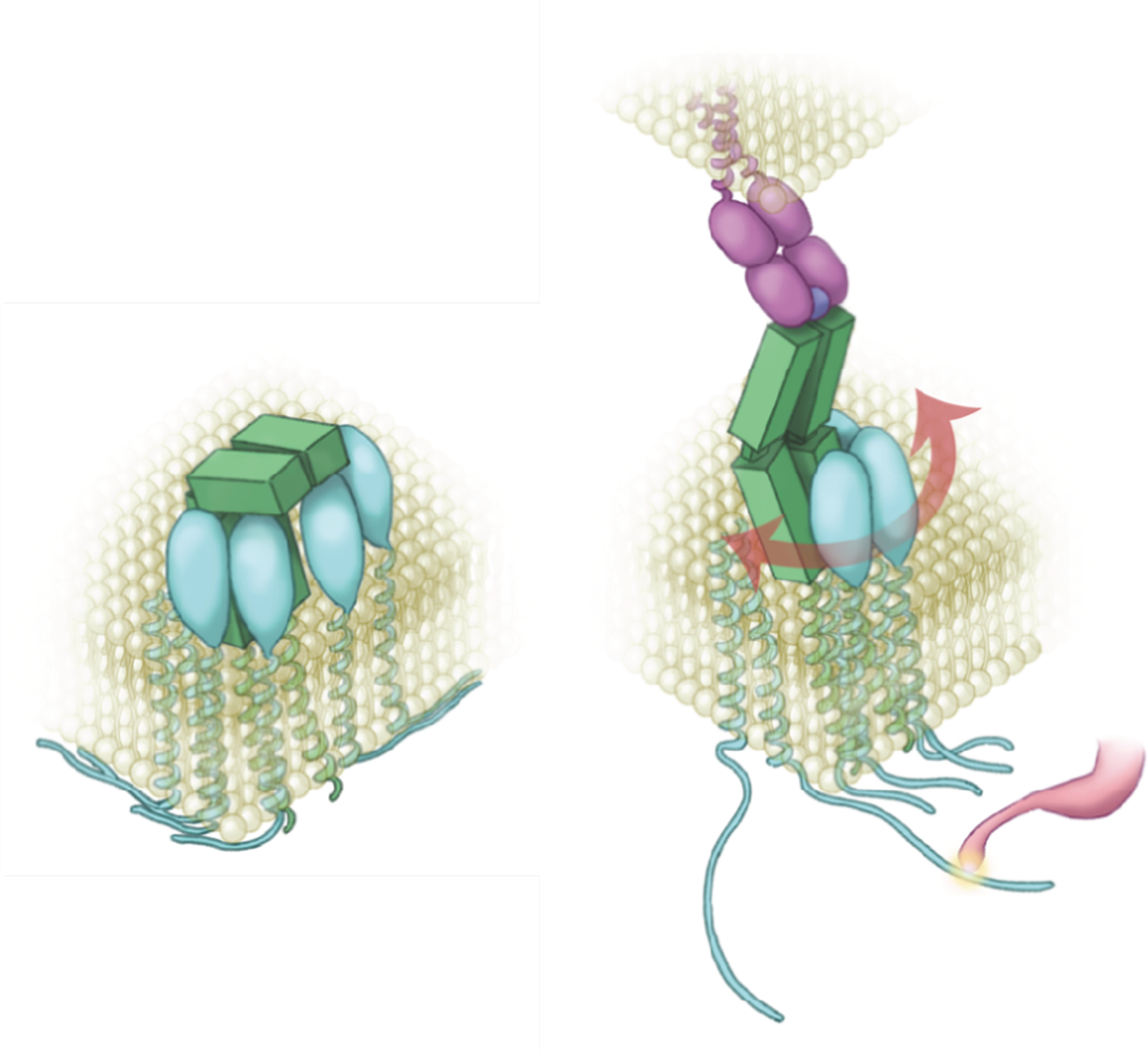
TCR-pMHC binding acts as a drawbridge for CD3 movement. An unbound TCR clamps the CD3 proteins, which prevents CD3 diffusion around the TCR in the membrane (*left*). Upon pMHC binding (*right*), the TCR becomes extended, which weakens binding to CD3 and allows CD3 diffusion around the TCR. Increased CD3 movement in the membrane facilitates the release of CD3 tails in the cytosol, where the ITAMS are accessible for phosphorylation by Lck.

### The TCRβ FG loop acts as a gatekeeper

Changes in the TCR have been observed upon TCR-pMHC binding that have been difficult to rationalize, because they occur far from the TCR-pMHC binding interface. Notably changes in the TCRɑ AB loop, the TCRβ H3 helix, and the TCRβ FG loop were observed by NMR^12, 20–22^ (Figure 6A). In our simulations, we observed an increase in the contacts between the TCRɑ AB loop and the TCRβ H3 helix with CD3 proteins upon TCR-pMHC binding (Figure 6B-D and Supplemental Figure S5). In unbound conditions the TCRɑ AB loop and the TCRβ H3 helix were out of reach of the CD3 proteins because they were bound by the TCR EC; however, upon TCR-pMHC binding, the CD3 proteins were released and moved to previously inaccessible areas of the TCR.

**Figure 6:**
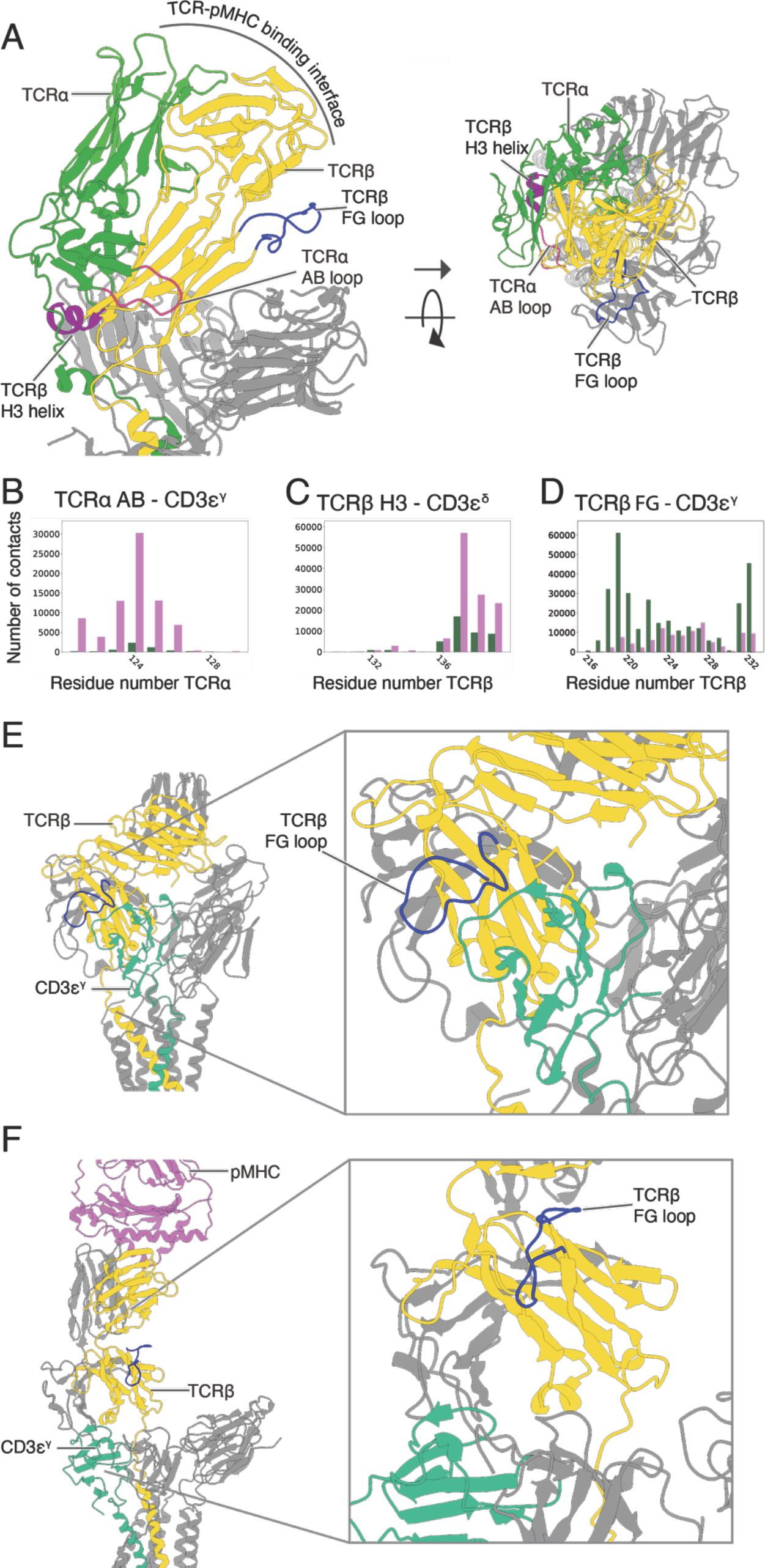
Contacts between TCRɑ AB loop, TCRβ H3 helix and the TCRβ FG loop and the CD3 proteins. (A) Side (*left*) and top view (*right*) of the TCR-CD3 cryo-EM structure. The TCRɑ AB loop, TCRβ H3 helix and the TCRβ FG loop are located far from the TCR-pMHC binding interface and are out of reach of the CD3 proteins. The CD3 proteins with extracellular domains (CD3δ, CD3ε and CD3γ) are located at the opposite side of the TCRɑβ chains, while the CD3ζ chains lack an extracellular domain. Approximate TCR-pMHC binding face is indicated. TCRɑ in green with the AB loop in fuchsia, TCRβ is shown in yellow with the H3 helix in purple and the FG loop in blue, the CD3 protein chains are colored grey for clarity. Structure from PDB code 6JXR^15^. Cumulative contacts between different CD3 chains and the TCRα AB loop (B), TCRβ H3 helix (C), and the TCRβ FG loop (D) in the TCR-CD3 (green) and TCR-CD3-pMHC simulations (purple). Contacts are summed over all simulations and shown as a graded color, where each step in the gradient represents contacts from a single simulation. Residue numbers in the TCRα or TCRβ chains are shown on the x-axis. (E-F) Simulation snapshots illustrating the role of the TCRβ FG loop. In the TCR-CD3 simulations the FG loop blocked the diffusion of CD3ε^γ^ (panel E). Upon TCR-pMHC binding, the TCR elongated, and the FG loop moved up, allowing the diffusion of the CD3 proteins alongside TCRβ. TCRβ is shown in yellow with the FG loop in blue, CD3ε^γ^ in viridian green, and the pMHC in purple, other protein chains are colored grey for clarity.

In our simulations we observed that the TCRβ FG loop acted as a gatekeeper, only allowing the diffusion of the CD3 proteins when the TCR was bound by pMHC. The TCRβ FG loop plays an important role in T cell activation^21, 23, 24^. Moreover, changes in the TCRβ FG loop have been observed upon TCR-pMHC binding^8, 12^. In our simulations we observed that when the TCR was not bound by pMHC, the TCRβ FG loop interacted with CD3ε^γ^ and blocked its diffusion away from the TCRβ chain (Figure 6D-F). Upon TCR-pMHC binding, the TCRβ FG loop moved up and no longer interacted with the CD3 proteins, which were subsequently free to diffuse around the TCRβ. TCRɑ lacks an equivalent loop, which explains the asymmetry in the iso-occupancy maps. For a more detailed description about TCR-CD3 contacts we refer to the supplementary section “*TCRɑ AB loop, the TCRβ H3 helix, the TCRβ FG loop and the CD3ζζ chains”*.

### Critical role of the connecting peptide in TCR dynamics

If TCR triggering depends on TCR EC movement, a change in TCR chain flexibility would be expected to affect TCR activation. We identified a conserved GxxG motif in the TCRβ connecting peptide (CP) that represents a potential hinge (Figure 7). Whereas the TCRɑ CP is long and flexible, the TCRβ CP is short, which increases the importance of the two glycine residues in the hinge. Moreover, the TCRɑ and TCRβ CPs are attached to each other by a disulfide bond, which suggests that global changes in the TCR are mediated by these glycine residues. To test whether the CP in TCRβ indeed functions as a hinge, we examined double alanine (AA) or proline (PP) substitutions using atomistic molecular dynamics simulations. Atomistic simulations with the CHARMM36m forcefield^25^ of a TCR-CD3 complex in the absence of pMHC were used to assess the effect of the CP on the intrinsic flexibility of the TCR EC.

**Figure 7:**
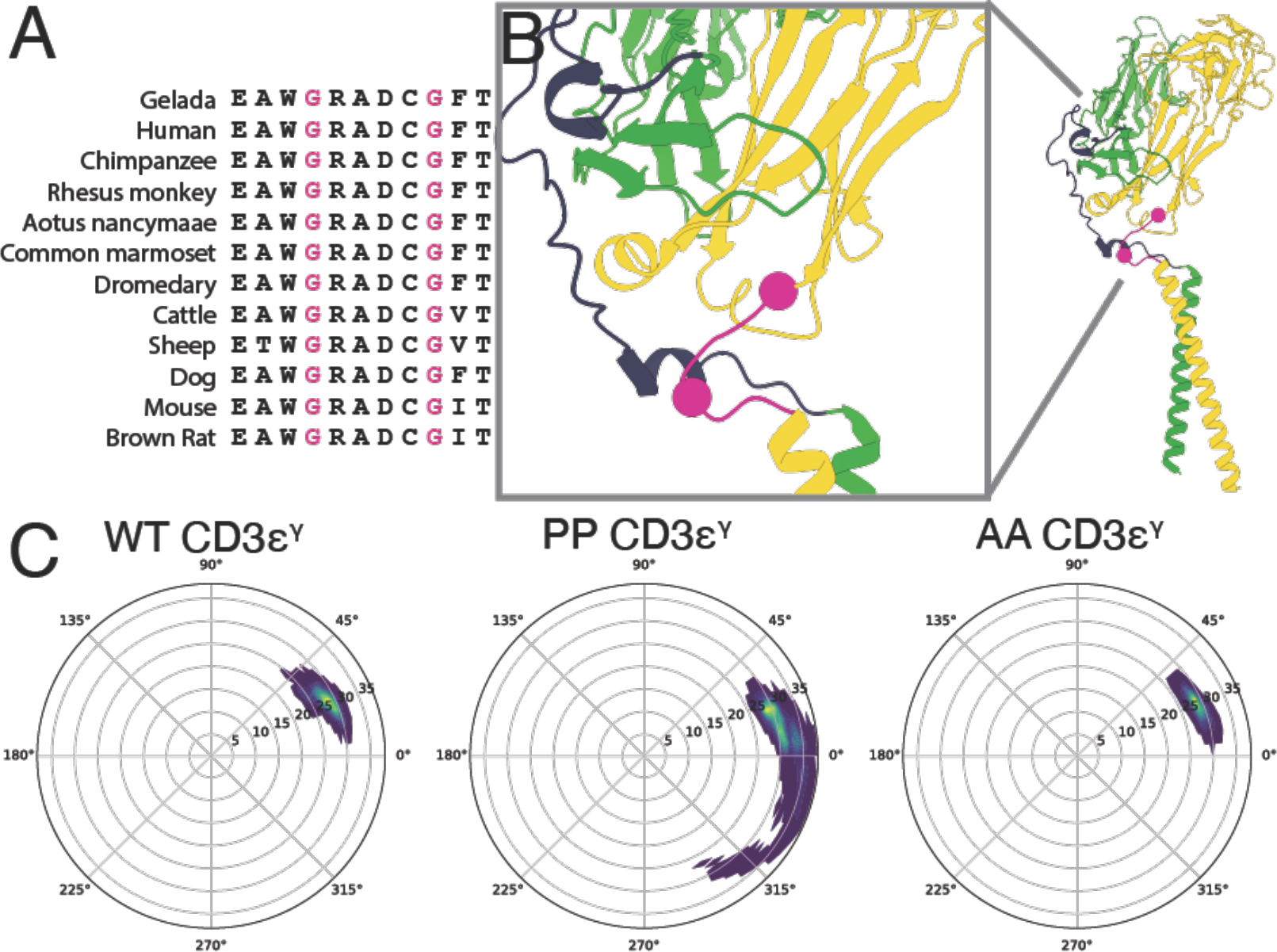
Hinge rigidification results in TCRs that are unable to retain the CD3 subunits. (A) Multiple sequence alignment of the TCRβ connecting peptide (CP) with the two conserved glycines in pink. (B) CP region of the TCR. The TCRɑ CP is longer than the TCRβ CP, but the TCRβ CP contains two flexible glycines. TCRɑ (green) with CP (blue), TCRβ (yellow) with CP (pink), and CP glycines (spheres) are shown. Structure from PDB code 6JXR^15^. (C) Atomistic molecular dynamics simulations for wild-type GG and “rigidified” (PP and AA) TCR mutants indicated that movement of CD3ε^γ^ (represented as densities in a 2D plane about the TCR) increased in the PP simulations compared to that in wild type.

The atomistic simulations showed that rigidification of the CP by glycine-to-proline mutations increased CD3ε dynamics, whereas a milder mutation of both glycines to alanines did not significantly affect CD3ε dynamics (Figure 7C, S6A). The mutations also changed the pattern of contacts between the TCR and CD3 chains. Both the proline and the alanine substitutions decreased the net number of interactions between the TCR variable region and CD3δε dimer as compared to wild type, but mutation to proline increased contacts between the TCRβ variable domain and CD3γ (Figure S6B). In line with the TCR-CD3 coarse-grained simulations, the wild-type and mutant TCRs adopted a bent conformation in the atomistic simulations with smaller EC-to-TM angles than that of the cryo-EM structure. The PP mutant adopted even smaller angles than the wild type, and, surprisingly, the bent conformations were decoupled from the CD3 subunits (Figure S6C-D). This suggested that the TCR tilt is not sufficient to describe the interactions between TCR and CD3. In the simulations of Pandey et al.^17^, the TCR EC not only varied its tilt, but also rotated. Indeed, calculation of the TCR EC rotation showed that the proline mutant had a skewed TCR EC – the PP mutant adopted rotation angles as low as ∼75°; in contrast, rotation angles lower than 100° were almost absent in the wild-type TCR (Figure S6E-F). The skewedness of the proline TCR was also apparent from the TCR and CD3 contacts– the wild-type TCR mostly contacted the CD3δε dimer, whereas the proline mutant interacted mostly with the CD3γε dimer located on the other side of the TCR (Figure S5B). The wild-type TCR restrained CD3δ via contacts between TCRɑ and the CD3δ FG loop, but the proline mutant could not reach CD3δ due to its skewed conformation (Figure S6G-H). These *in silico* mutation simulations were consistent with a model in which the CP functions as a hinge for the drawbridge, and that rigidification of the hinge by proline substitution locks the bridge in a skewed state in which the TCR can no longer properly restrain the CD3 chains, resulting in their increased mobility.

### T cells harboring rigidified TCRs are hypersensitive

The importance of CP flexibility for TCR activation was next investigated experimentally. Variants of the 3A9 TCR^26^ were created in which the two glycine residues in the TCRβ CP were mutated to either proline (PP) or alanine residues (AA). NFAT-GFP reporter cell lines^27^ were transfected with one of three different TCRs: wild type (GG), PP mutant, or AA mutant. The NFAT-GFP cells produce GFP upon T cell activation, which is used as a reporter for TCR triggering^26, 27^. The reporter cells were co-cultured with antigen presenting cells and different concentrations of HEL peptide. After 2, 4 or 24 h of co-culture, T cell activity was measured by determining the percentage of GFP-expressing T cells by FACS (Figure 8A-D, Supplemental Figures S7-S8). Proline substitution resulted in a higher percentage of activated T cells, whereas alanine substitution resulted in only a slightly higher percentage of active T cells compared to that in wild type. The effect was especially clear after 2 h but decreased at later timepoints.

**Figure 8:**
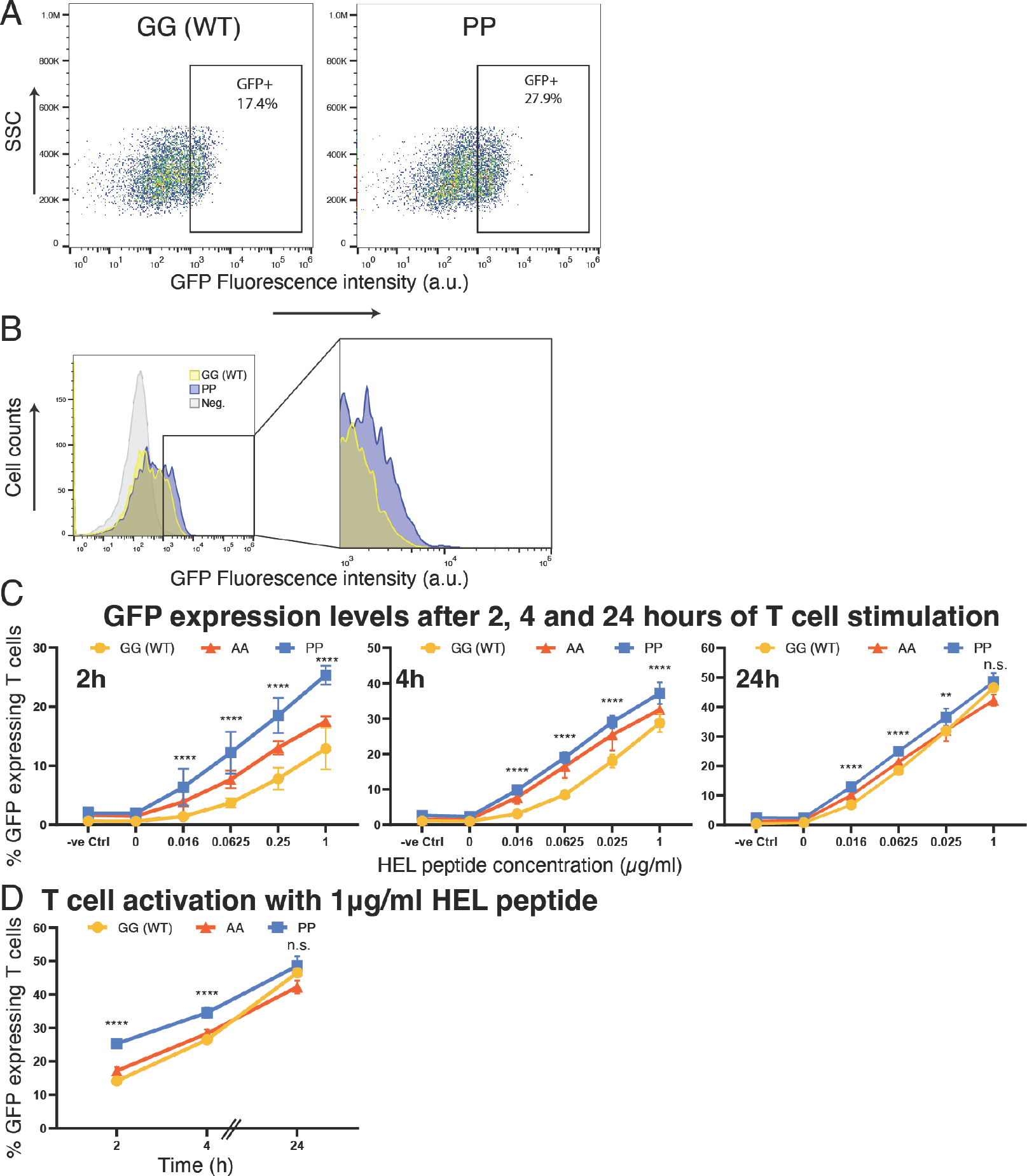
Hinge rigidification results in hyperresponsive T cells. NFAT reporter cells were transfected with GG, AA, or PP mutant TCRs. Cells with the PP TCRs, but not GG or AA TCRs, were hyperresponsive to pMHC stimulation after 2 h, but the difference between mutant and wild type cells decreased after 4 h and 24 h. Flow cytometry dot plots (A) and histogram (B) identifying GFP-positive T cells after co-culturing with APC and 1 µg/ml HEL peptide for 2 h. Results of a representative experiment are shown. (C) Percentage of GFP expressing T cells after 2, 4 or 24 h of co-culture. Note that the y-axis varies between the plots. Data are represented as mean ± stdev for all replicates from 2-3 independent experiments. (D) Percentage of GFP-positive 3A9 T cells after being stimulated with 1 μg/ml HEL peptide over different time points (2, 4, 24 h) Data are represented as mean ± stdev for 3 replicates from one representative experiment. n.s.: non-significance, ****: P <0.0001 for PP compared to wild type (GG), **: P <0.01 for PP compared to wild type (GG).

These experiments showed that reporter cells bearing the mutant TCR were more easily activated, which suggests that rigidification of the CP by mutation to proline, and to a lesser extent to alanine, resulted in TCR hypersensitivity. This is in line with the atomistic simulations in which the proline mutant was less able to couple to the CD3 proteins due to rigidification of the CP and skewing of the TCR EC. These experiments provide support for the hypothesis that the TCR CP functions as a hinge and underlie the importance of TCR EC movement and CD3 mobility for TCR triggering.

## Discussion

The mechanism of TCR triggering in which TCR-pMHC binding is signaled across the plasma membrane, resulting in phosphorylation of ITAMs on cytoplasmic CD3 tails, is still not fully resolved. It has been difficult to conceptualize how TCR-pMHC binding results in ITAM phosphorylation: the TCRɑβ chains lack a proper cytoplasmic domain and the CD3 proteins are too short to interact with the pMHC. In this study, the behavior of the TCR-CD3-pMHC complex was investigated using extensive multi-scale simulations, resulting in a model in which TCR-pMHC binding allows dynamic movement of the CD3 proteins.

Coarse-grained simulations showed that the TCR adopts a bent conformation when not bound to pMHCs. In this bound state, the TCR EC binds to the CD3 proteins, restricting their movement. The TCRβ FG loop hereby acts as a gatekeeper, preventing accidental TCR triggering by CD3 diffusion when the TCR is unbound. TCR-pMHC binding requires the TCR to elongate, breaking its interactions with the CD3 proteins and allowing them to diffuse freely. The simulations also indicated the importance of the CP, a flexible region linking the TCR’s transmembrane region with its extracellular domain, for the TCR to adopt the necessary conformational changes. The TCRα CP is longer, while the TCRβ CP is much shorter but contains two flexible glycine residues. Together the CPs allow the TCR-pMHC to act like a “drawbridge”, which either blocks the CD3 proteins or lets them pass freely.

Atomistic simulations and experiments provided further support for the crucial role of the TCR CP. The simulations showed that TCR-CD3 complexes in which the two glycines in the CP were replaced with prolines resulted in a skewed TCR that was less effective in retaining the CD3 proteins. This was confirmed by experiments using reporter cells transfected with either wild-type or mutant TCRs. Activation of reporter cells with the proline mutant TCR was found to be more efficient than wild type, suggesting that CD3 mobility is a key step in TCR triggering.

The drawbridge model provides a solution to how TCR-pMHC binding is signaled to the CD3 proteins but does not reveal how increased CD3 mobility upon TCR-pMHC binding results in ITAM phosphorylation. The predominant theory is that the ITAMs are sequestered in the membrane, but that changes in the lipid microenvironment upon TCR-pMHC binding results in their release from the membrane and their exposure to Lck^2, 14, 28^. It is reasonable to hypothesize that, in the resting state, when the TCR is not bound to pMHC, the CD3 cytosolic tails are sequestered in lipid rafts. Increased CD3 dynamics upon TCR-pMHC binding would likely disrupt these lipid rafts by inducing a disordered phase, dispersing the lipids. If ITAMs in particular associate with certain lipids in these rafts, such as charged lipids, disruption of lipid rafts could result in the release of ITAMs. This hypothesis is supported by two experimental observations: first, lipid rafts are known to be present in the T cell membrane and are involved in T cell signaling^29–32^; second, cholesterol, which induces liquid-ordered phases and is an important component of lipid rafts, has been shown to decrease TCR triggering^28, 33, 34^. Further work will be needed to elucidate the detailed implications of CD3 dynamics on ITAM phosphorylation, but the overall notion that triggering is associated with greater CD3 dynamics appears sound.

The drawbridge model posits that the TCR is flexible and adopts a bent conformation when not bound to a pMHC. However, this conformation has not yet been directly observed experimentally; rather, in the three reported single particle cryo-EM structures of the TCR-CD3 complex, the TCR exists in an extended conformation, similar to that observed in our TCR-CD3-pMHC simulations^15, 18, 28^. One possible explanation for the structural homogeneity observed in the reported single particle cryo-EM structures is that, because of the experimental procedures used in the preparation of the TCR-CD3 complex for structure determination, the physiological environment of the cell membrane had to be removed. In addition, it was also required to stabilize the complex by the use of chemical crosslinking, introduction of antibodies or the use of specialized cryo-grids in order to arrive at a consensus structure^35^. The study by Dong and co-workers^15^, for example, employed 0.1% glutaraldehyde for cross linking transmembrane helices while that of Sušac and co-workers used graphene coated grids and Fabs for TCR stabilization^18^. The absence of a native membrane is especially relevant since movement of the TCR-EC was found to be coupled to changes in the transmembrane helices in the previous work by Pandey and co-workers^17^. Furthermore, in TCR-CD3 MD simulations reported to date, TCR bending was observed consistently, despite the use of three different forcefields – Amber99SB-ILDN^17^, and CHARMM36m and the Martini 3 forcefields, in this work. There is, moreover, experimental support for a reversible conformational change in the TCR from single molecule studies of isolated cells using optical tweezers^36^. Similar work on the pre-TCR resulted in a model in which the pre-TCR EC moves due to the scanning motion of the APC^37^. Consistently, changes in chemical shift were observed not only in constant regions, but also in the TCRɑβ variable regions upon titration with CD3γε and CD3δε^38^, providing support for the idea that the TCRɑβ variable region interacts dynamically with CD3 proteins.

There is also experimental support for the drawbridge model in the literature. Long-range perturbations have been observed in the TCRβ FG loop, TCRα AB loop, and TCRβ H3 helix upon TCR-pMHC binding using a range of techniques, including NMR^20–22, 38^, optical tweezers^8^ and fluorescence measurements^39^. These studies have revealed structural changes in the TCR far from the binding interface upon TCR-pMHC binding that have been difficult to rationalize. Our model argues that the long-range interactions are due to changes in the TCR-CD3 interactions. In the unbound state, the CD3 proteins are bound by the TCR EC, but upon TCR-pMHC binding the TCR elongates and the CD3s diffuse around the TCR. The CD3 proteins are thus able to interact with regions like the TCRα AB loop and TCRβ H3 helix of the TCR, which are not accessible in the unbound state. The TCRβ FG loop has thereby a special role as a gatekeeper, blocking the movement of the CD3 proteins by interacting with CD3ε^γ^ in the unbound state, but moving out of the way after TCR-pMHC binding, and thus preventing inadvertent TCR triggering. Additional support comes from hydrogen-deuterium exchange experiments, which point to a rigidification of the TCR upon TCR-pMHC binding^40^, which is consistent with the narrower angle distribution of the pMHC bound TCR in our simulations (Figure 4B). Furthermore, experimental studies have shown that a decrease in TCR-CD3 interactions occurs upon TCR-pMHC binding^13^ and specifically between the TCR and CD3ζ^14^. Consistently, our simulations also show a decrease in TCR-CD3ζ interactions upon TCR-pMHC binding (Figure S3). This decrease in TCR-CD3 association is in line with the increased mobility of the CD3 and an elongated TCR that cannot interact with the CD3 upon TCR-pMHC binding. Lastly, molecular docking studies utilizing NMR data^38^ resulted in a model in which the CD3δε and CD3γε dimers were positioned on opposite sides of the TCRαβ complex. This model differs qualitatively from the recent cryo-EM structures^15, 18^, providing support for a model based on CD3 mobility and structural heterogeneity.

The drawbridge model is consistent with recent findings that monomeric TCRs can be triggered^41^ and integrates well with the modern framework of TCR triggering, such as the mechanosensing^7, 42^ and kinetic proofreading^10^. The initial encounter between TCR and pMHC is described by the mechanosensing model, which posits that agonistic pMHCs are distinguished through the formation of catch-bonds, special bonds that withstand the shear stress created by the scanning motion of the T cell over the APC^4, 43^. The drawbridge model expands on this by proposing that these catch-bonds must also be strong enough to maintain the TCR in the elongated state. The kinetic proofreading model enhances the specificity of T cell activation by imposing a temporal requirement, such that only TCR-pMHC interactions that persist for a sufficient amount of time result in T cell activation^10, 44, 45^. This is achieved via a series of reversible biochemical steps, and the elongation of the drawbridge can be seen as part of this process. Successful TCR triggering can only occur when the TCR remains in an elongated state for long enough to allow for the continuous diffusion of the CD3 proteins, which ultimately depends on the stability of the complex formed between the TCR and pMHC. Our molecular dynamics simulations thus link the upstream formation of catch-bonds with the downstream kinetic proofreading process, providing a more comprehensive understanding of TCR triggering.

In conclusion, a wide range of evidence supports a mechanism in which TCR-pMHC binding is coupled to the dynamics of the CD3 EC domains, which is propagated across the plasma membrane. Further work is needed to confirm the flexibility of the TCR headgroup and to investigate how CD3 diffusion results in ITAM phosphorylation. Another interesting direction is the exploitation of this model for manipulation of T cell activation on demand. In this context, it will be interesting to revisit the molecular mechanism of anti-CD3 antibodies, which are used to control T cell activation both in the laboratory and in the clinic^46–48^.

## Acknowledgments

Simulations were performed using the Research Center for Computational Science, Okazaki, Japan (Projects: 20-IMS-C116, 21-IMS-C116 and 22-IMS-C116) and the TSUBAME3.0 supercomputer at Tokyo Institute of Technology. This work was supported by Japan Agency for Medical Research and Development (AMED), Platform Project for Supporting Drug Discovery and Life Science Research (Basis for Supporting Innovative Drug Discovery and Life Science Research) under JP22ama121025. The authors wish to thank Misaki Ouchida for creating the illustration explaining the drawbridge model and Robert Mallis for the careful reading of the manuscript.

## Author contributions

Conceptualization and supervision were done by F.J.V.E. and D.M.S.; *in silico* simulations were performed by F.J.V.E. and analyzed by F.J.V.E. and D.M.S. All authors contributed to the *in vitro* experimental design and analysis, but *in vitro* experiments were performed by A.A.S., M.A.L.C. and A.M. Original draft was written by F.J.V.E., D.M.S., A.A.S., M.A.L.C and A.M. All authors reviewed the manuscript.

## Methods

### Computational methods

Our aim was to examine a conventional TCR-MHC complex, but the extracellular part of the TCR electron densities of the De Dong et al. structure were fitted to the coordinates of an atypical natural killer TCR. Therefore, a chimeric structure was created by fusing the structure of De Dong et al. (PDB ID 6jxr)^15^ with the structure of a conventional TCR from Newell et al. (PDB ID 3qiu)^19^. For the CD3 proteins and the transmembrane part of the TCR, the structure of De Dong was used. The ITAM fragments were not present in the structure by De Dong et al. and were modeled according to the murine sequence (see below). The structure by Newell et al. served as the basis for the MHC and the extracellular parts of the TCR. Please note that the structure by De Dong et al. is human, but the Newell et al. structure is in itself already a chimeric structure in which the constant domains are human and the variable domains murine. The ITAMs were modelled according to the murine sequences.

### MHC

The structure from Newell et al. lacks MHC transmembrane helices, which were modeled using HHpred^49^ and Pymol^50^. To obtain a realistic orientation of the two MHC transmembrane helices, a 10-µs coarse-grained molecular dynamics simulation of the two helices embedded in a POPC bilayer composed out of 221 lipids was performed with the martini 3 force field (version v.3.0.4.26). The modeled atomistic helices were subsequently fitted on the simulated structures and connected to the soluble part of the MHC from Newell et al. using Modloop^51^. The epitope was restrained to the MHC using a number of harmonic potentials (Gromacs bondtype 6), see Table 4.

### TCR

For the simulation, a fusion of the De Dong et al. and Newell et al. structures was created. From the structure of De Dong, the CD3s were used, and the transmembrane segments of the α and β chains, which correspond to residue numbers 224-273 for the α chain and residues 264-308 for the β chain. The soluble segments of the TCRαβ chains were taken from the structure of Newell et al., for the α chain residues 2-199 and for the β chain residues 2-244, the residue numbering is according to the respective pdb files. The structures were fused using Pymol^50^ and Modloop^51^. Please note that the TCR in the Newell structure has an engineered disulfide bond between TCRα Cys156 and TCRβ to improve the stability of the structure for crystallization^19^.

Part of the missing and incomplete TCR and MHC residues were reconstructed using Spanner^52^ and Pymol^50^ (Table 3). Residues included in the simulation are shown in Tables 1 and 2. This structure forms the basis for the fine grained (FG) and the coarse grained (CG) simulations. For the CG but not the FG simulations, ITAMs (immunoreceptor tyrosine-based activation motif) were modeled using Pymol. The TCR was subsequently coarse grained (see below) and embedded in a membrane (see below). A set of equilibration simulations was performed to equilibrate the ITAM chains. For the FG simulations, the system was prepared using CHARMM-GUI^53^.

**Table 1:**
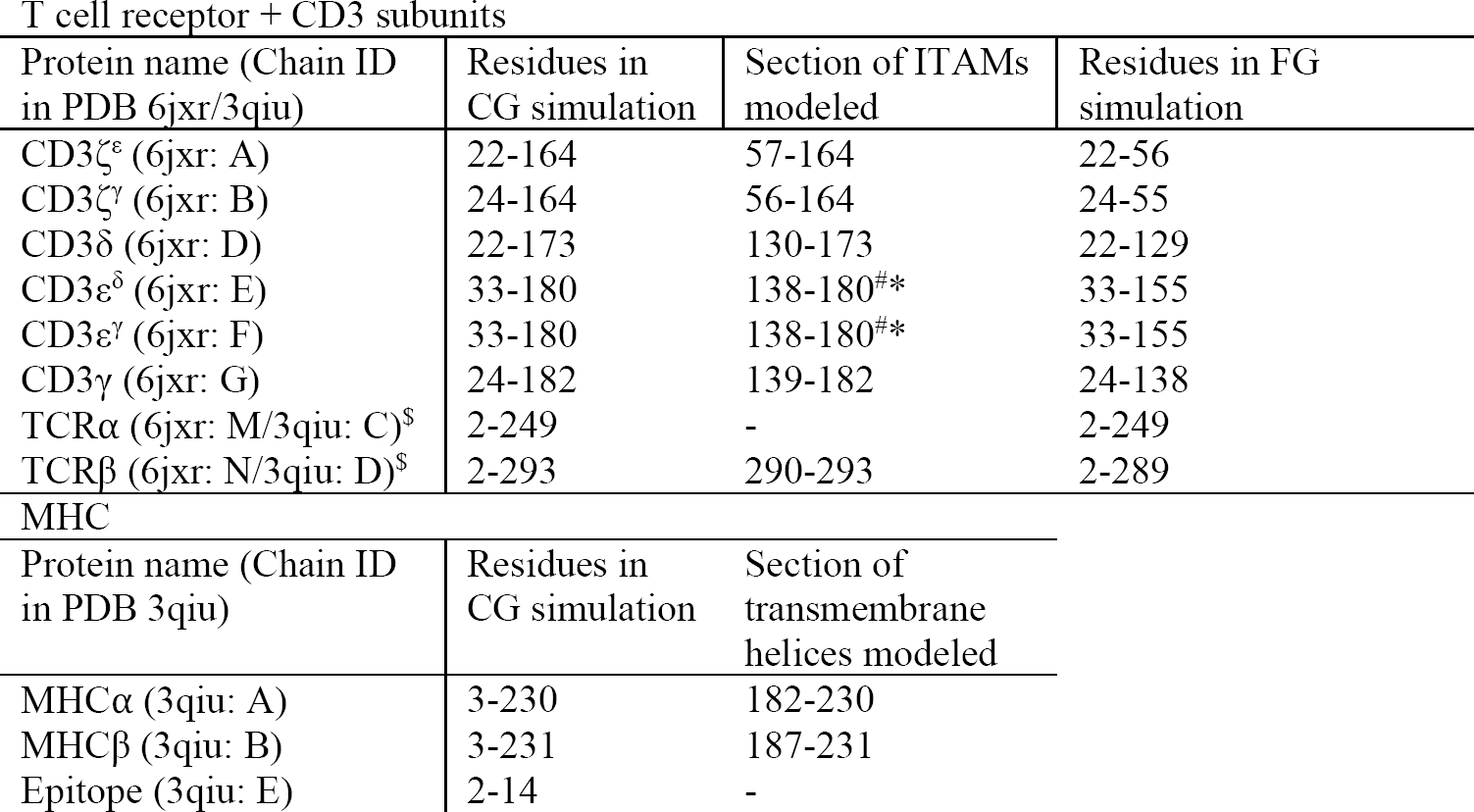
Residues included in the simulation. In the ITAM column, the numbers between the brackets refer to the uniprot numbering ^#^The CD3ε ITAM sequence length differs between mouse and human, the numbers refer to mouse sequence. *Residues 181-189 of the CD3ε chains (TCR E and TCR F) were omitted during modeling and not included in the simulation. ^$^Numbering scheme of 3qiu is used, chain C for TCR ⍺ and chain D for TCRβ.

**Table 2:**
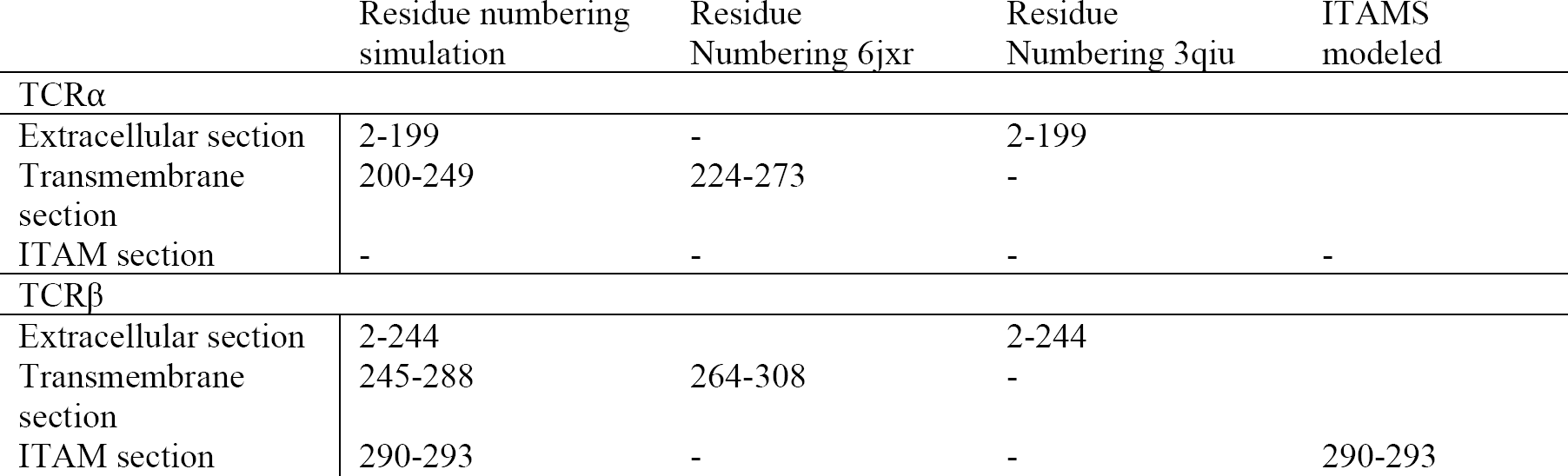
Residues included in the simulation. In the ITAM column, the numbers between the brackets refer to the uniprot numbering ^#^The CD3ε ITAM sequence length differs between mouse and human; the numbers refer to the mouse sequence. *Residues 181-189 of the CD3ε^δ^ and CD3ε^γ^ chains (TCR E and TCR F) were omitted during modeling and not included in the simulation. ^$^Numbering scheme of 3qiu is used, chain C for TCR⍺ and chain D for TCRβ.

**Table 3:** TCR and MHC residues used for reconstruction. Unless indicated, residues were reconstructed using Spanner^52^.

| Protein | PDB ID | Chain | Residues |
| --- | --- | --- | --- |
| MHCβ | 3qiu | B | 105-113,134-135,165-170 |
| TCRα | 3qiu | C | 145-146, 175-179, 187-199 |
| TCRβ | 3qiu | D | 183-184, 219-220, 244 |
| CD3ζ | 6jxr | A | 57 <sup>*</sup>modeled with Pymol |
| CD3ε | 6jxr | F | 156 <sup>*</sup>modeled with Pymol |

**Table 4:** Harmonic potentials used to restrain the epitope in the MHC.

| MHC<br>chain | MHC<br>residue<br>number | MHC<br>residue | MHC<br>bead | Epitope<br>residue<br>number | Epitope<br>residue | Epitope<br>bead | Equilibrium<br>distance<br>(nm) | Force<br>constant<br>(kJ mol<sup>-1</sup><br>nm<sup>-2</sup>) |
| --- | --- | --- | --- | --- | --- | --- | --- | --- |
| A | 52 | ALA | BB | 2 | ALA | BB | 0.460 | 200 |
| B | 86 | PHE | SC2 | 5 | ILE | SC2 | 0.462 | 200 |
| A | 54 | PHE | SC2 | 5 | ILE | SC2 | 0.490 | 200 |
| B | 11 | CYS | SC1 | 10 | GLN | SC1 | 0.517 | 200 |
| B | 30 | TYR | SC4 | 11 | ALA | SC4 | 0.490 | 200 |
| A | 69 | ASN | SC1 | 12 | THR | SC1 | 0.507 | 200 |

**Table 5:**
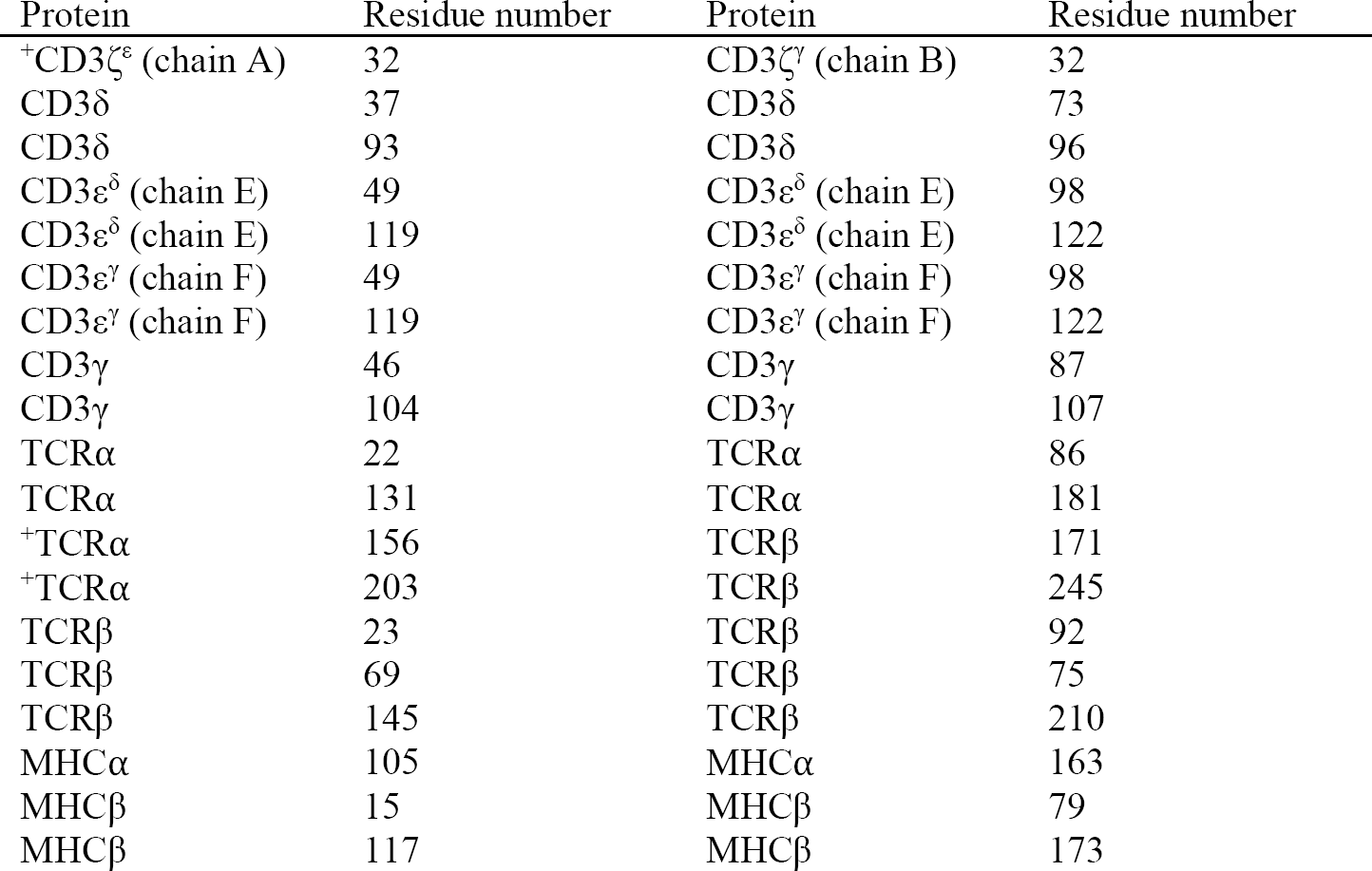
Disulfide bonds present in the simulation. ^+^Interchain disulfide bond are modeled by a harmonic potential.

### Positioning of TCR and MHC

TCR and pMHC were carefully positioned on top of each other in preparatory 1.5 µs simulations. The TCR and MHC were embedded in opposing membrane slabs, which were separated by ∼21 nm (GL1-GL1 bead distance) (Figure 1). A set of distance restraints based upon the Newell structure (PDB: 3qiu)^19^ was defined between the TCR and pMHC (Table 6). During the 1.5 µs docking simulation, a low force constant of 1 kJ mol^−1^ nm^−2^ was used to prevent any deformations. During the docking simulation, the TCR and pMHC were slowly pulled together by distance restraints resulting in a bound TCR-CD3-pMHC structure (Figure S1). In the production simulations, a higher force constant of 200 kJ mol^−1^ nm^−2^ was used to keep the pMHC properly bound to the TCR. In order to allow solvent flow between the two membrane compartments, which in turn allows the bilayers to move, a pore was created in the bilayer containing the MHC. The pore was created using two cylinder shaped inverted flat bottom position restraints (function type 2, shape *g* = 8 in Gromacs). The first restraint, which was used to create the actual pore, acted on POPC beads with a force constant of 1000 kJ mol^−1^ nm^−2^ and a radius of 2 nm. The second restraint with a force constant of 2000 kJ mol^−1^ nm^−2^ and a radius of 4 nm acting on the MHC backbone beads was used to prevent MHC TM diffusion into the pore. The preparatory simulations were performed for each of the one hundred TCR-CD3-pMHC simulations independently.

**Table 6:** List of distance restraints between the TCR and the pMHC. During the docking simulations, the force constant was set to 1 kJ mol^−1^ nm^−2^. A piecewise linear/harmonic restraint was used (bond type 10 in Gromacs) to keep the TCR and the pMHC together. The potential is quadratic for rij < r0, 0 for r0 ≤ rij < r1, quadratic for r1 ≤ rij < r2, and linear for r2 ≤ rij.

| MHC chain | MHC residue number<br>r | MHC residue | MHC bead | TCR chain | TCR residue number | TCR residue | TCR bead | $r_0$ (nm) | $r_1$ (nm) | $r_2$ (nm) | Force constant (kJ mol<sup>-1</sup> nm<sup>-2</sup>) |
| --- | --- | --- | --- | --- | --- | --- | --- | --- | --- | --- | --- |
| MHCβ | 69 | GLU | BB | TCRα | 49 | ALA | BB | 0.593 | 0.693 | 3.093 | 200 |
| MHCβ | 77 | THR | BB | TCRα | 29 | ARG | BB | 0.482 | 0.582 | 3.682 | 200 |
| MHCβ | 84 | GLU | BB | TCRα | 65 | LYS | BB | 1.102 | 1.202 | 4.302 | 200 |
| MHCα | 57 | GLN | BB | TCRβ | 57 | GLN | BB | 0.788 | 0.888 | 3.988 | 200 |
| MHCα | 61 | ALA | BB | TCRβ | 33 | PHE | BB | 1.211 | 1.311 | 4.411 | 200 |
| MHCα | 75 | GLU | BB | TCRβ | 53 | GLU | BB | 1.800 | 1.900 | 5.000 | 200 |
| MHCε | 6 | ALA | BB | TCRα | 92 | SER | BB | 0.320 | 0.480 | 3.520 | 200 |
| MHCε | 11 | ALA | BB | TCRβ | 98 | ASN | BB | 0.429 | 0.529 | 3.629 | 200 |
| MHCα | 61 | ALA | BB | TCRβ | 55 | LEU | BB | 0.365 | 0.465 | 3.565 | 200 |
| MHCα | 64 | ALA | BB | TCRβ | 55 | LEU | BB | 0.560 | 0.660 | 3.760 | 200 |
| MHCα | 57 | GLN | BB | TCRβ | 55 | LEU | BB | 0.739 | 0.839 | 3.939 | 200 |
| MHCα | 64 | ALA | BB | TCRβ | 53 | GLU | BB | 0.649 | 0.749 | 3.849 | 200 |
| MHCα | 57 | GLN | BB | TCRβ | 53 | GLU | BB | 1.327 | 1.447 | 4.547 | 200 |
| MHCα | 61 | ALA | BB | TCRβ | 57 | GLN | BB | 0.917 | 1.067 | 4.167 | 200 |
| MHCα | 64 | ALA | BB | TCRβ | 57 | GLN | BB | 1.231 | 1.331 | 4.431 | 200 |
| MHCα | 57 | GLN | BB | TCRβ | 33 | PHE | BB | 1.421 | 1.521 | 4.621 | 200 |
| MHCα | 64 | ALA | BB | TCRβ | 33 | PHE | BB | 1.441 | 1.541 | 4.641 | 200 |
| MHCα | 64 | ALA | BB | TCRβ | 49 | PHE | BB | 0.899 | 0.999 | 4.099 | 200 |
| MHCα | 65 | VAL | BB | TCRβ | 49 | PHE | BB | 0.916 | 1.016 | 5.116 | 200 |
| MHCα | 61 | ALA | BB | TCRβ | 49 | PHE | BB | 0.843 | 0.943 | 5.043 | 200 |
| MHCβ | 73 | ALA | BB | TCRα | 49 | ALA | BB | 0.528 | 0.728 | 1.828 | 200 |
| MHCβ | 66 | GLU | BB | TCRα | 49 | ALA | BB | 0.706 | 0.806 | 2.006 | 200 |
| MHCβ | 73 | ALA | BB | TCRα | 29 | ARG | BB | 0.664 | 0.764 | 1.964 | 200 |
| MHCβ | 74 | GLU | BB | TCRα | 29 | ARG | BB | 0.625 | 0.725 | 3.825 | 200 |
| MHCβ | 76 | ASP | BB | TCRα | 65 | LYS | BB | 0.643 | 0.743 | 3.602 | 200 |
| MHCβ | 73 | ALA | BB | TCRα | 65 | LYS | BB | 0.930 | 1.030 | 4.130 | 200 |

### Coarse graining and run parameters

Simulations were performed with the Martini 3 force field, version v.3.0.4.26^54^ in combination with a Gō-like model^55, 56^. Coarse-grained topologies were created using Martinize2^57^. Simulations were performed with Gromacs, versions 2018-2022^58^ with a 20 fs timestep and the ‘New-RF’ parameter set for the Martini force field^59^ with the modification that the Berendsen barostat was used. Temperature was kept at 310 K using the V-rescale thermostat^60^ using a coupling constant of τ_t_ = 1 ps. Pressure was semi-isotropically coupled to an external bath of p = 1 bar with a coupling constant of τ_p_ = 4 ps and a compressibility of χ = 3 ×10^−4^ bar^−1^ using the Berendsen barostat^61^.

### Fine-grained simulations and run parameters

For the FG simulations the structure of the chimeric TCR, excluding the ITAMs, was processed with CHARMM-GUI^53, 62^ and simulations were carried out with the CHARMM36m forcefield^25^ and Gromacs version 2021.2^58^. Simulations were performed with the parameters provided by CHARMM-GUI. In short, a time step of 2 fs was used, and the electrostatics were calculated using particle-mesh Ewald with a cutoff of 1.2 nm^63, 64^. Van der Waals interactions were calculated using a cutoff of 1.0 nm and the forces were switched to zero between 1-1.2 nm. Temperature was kept at 310 K using the Nose Hoover thermostat^65, 66^ using a coupling constant τ_t_ = 1 ps. Pressure was semiisotropically kept at 1 bar using a coupling constant of τ_p_ = 5 ps and a compressibility of χ = 4.5 ×10^−5^ bar^−1^ using the Parrinello-Rahman barostat^67^.

The TCR-CD3 systems were embedded in bilayers composed of 562 POPC lipids. The systems were solvated in water and 154 mM NaCl and 19 Na^+^ counterions to neutralize the systems. Due to the stochastic nature of system construction, the number of solvent molecules differed slightly between the systems. The wild-type system was solvated with 79097 and 222 NaCl ion pairs, the proline system with 78837 H_2_O molecules and 221 NaCl ion pairs and the alanine system with 78764 H_2_O and 221 NaCl.

### Simulation systems

The protein systems were embedded in POPC bilayers using the Insane script^68^. The TCR was embedded in a POPC bilayer of 1279 lipids (651 lipids in the extracellular leaflet and 628 lipids in the intracellular leaflet). The pMHC was embedded in a POPC bilayer of 1278 lipids (640 lipids in the extracellular leaflet and 638 lipids in the intracellular leaflet). The TCR-only system was solvated in 90282 water beads (representing 361128 atomistic waters) and 1018 Na^+^Cl^−^ beads were added (around 154 mM) and an additional 6 Cl^−^ ions to neutralize the system. The TCR-CD3-pMHC system was solvated in 95973 water beads (representing 383892 atomistic waters) and 1079 Na^+^Cl^−^ beads were added (around 154 mM) and an additional 12 Na^+^ ions to neutralize the system. One hundred independent 10-µs production simulations were performed for both the TCR-CD3 and TCR-CD3-pMHC systems. Note that each TCR-CD3-pMHC production run was preceded by a 1.5 µs preparatory simulation in which the TCR was slowly docked to the pMHC.

### Analysis

The first 1 µs of each coarse-grained production simulation was discarded as equilibration, and the remaining (9 µs) trajectories were analyzed every 10 ns. The first 250 ns of the atomistic simulations were discarded as equilibration and the remaining (2.25 µs) trajectories were analyzed every 1 ns. The TCR tilt, defined as the angle between the TCRβ EC and TM was calculated for simulation systems between the backbone beads (coarse-grained systems) or Cɑ atoms (atomistic simulations) of asparagine 97 serine 249 and valine 281 of the TCRβ chain. The tilt of the TCR in the cryo-EM structure was calculated between the Cɑ atoms of arginine 114, serine 268 and valine 300. Note that serine 249/268 and valine 281/300 are the corresponding residues in the simulation systems and cryo-EM structure, arginine 114 was chosen for its proximity to asparagine when superimposing the simulated structure and the cryo-EM structure. The rotation of the TCR EC was calculated as the angle between two planes. Plane 1 was defined by the Cɑ atoms used for the tilt TCR EC tilt calculation: asparagine 97 serine 249 and valine 281 of the TCR β chain. Plane 2 was defined by the Cɑ atoms of TCRβ asparagine 97, TCRβ glycine 14 and TCRɑ serine 18. Iso-occupancy maps were calculated using the Volmap tool in VMD. For the iso-occupancy maps both the backbone and the sidechain beads were used. Grid spacing was set to 2 Å and counting of grid occupancy was performed after fitting the trajectories on the backbone beads of the TCR chains. Contacts between the TCR and CD3 chains were calculated with in-house python scripts making use of the MDAnalysis library^69, 70^. For both coarse-grained and atomistic simulations, beads or atoms were considered in contact when their distance was < 6 Å, and residue contacts were calculated by summing all bead or atom contacts. The following residues were considered to be part of the constant, variable, transmembrane and cytosolic domains for the TCR chains: TCRα variable: 1-106, constant: 107-198, transmembrane: 199-246, cytosolic: 247-249. TCRβ variable: 1-117, constant: 118-245, transmembrane: 246-285, cytosolic: 286-293. The polar plots were calculated by setting the center of geometry of the TCRβ as the origin, and by representing the CD3ε^γ^ and CD3ε^δ^ chains by their centers of geometry. Centers of geometry were calculated using both backbone and side-chain beads (coarse-grained simulations) or atoms (atomistic simulations). Trajectories were viewed using VMD and ChimeraX and pictures were rendered with ChimeraX^71–73^.

## Experimental methods

### Objective

In this study, we investigated two glycines (G240 and G245) in the TCRβ connecting peptide. We generated three NFAT-GFP reporter T cells^27^ that express 3A9 TCR as model systems. One expressed a wild-type (GG) 3A9 TCR connecting peptide and the other two expressed mutated hinge regions at G240 and G245 with either alanine (AA) or proline (PP). T cells were co-incubated with antigen presenting cells and cognate peptide, and cell activation was evaluated by measuring GFP expression using flow cytometry.

### Peptide

Hen egg lysozyme (HEL) (46-61) peptide specific for 3A9 hybridoma and was obtained from GeneScript (#RP11736) at a purity of more than 95%. Its amino acid sequence is NTDGSTDYGILQINSR. It was reconstituted in sterilized water at a concentration of 1000 μg/ml and stored at -20°C.

### Cell Lines

Phoenix cells, obtained from ATCC, were cultured in complete DMEM media [cDMEM; DMEM (FUJIFILM Wako, Osaka, Japan) supplemented by 10% heat-inactivated fetal bovine serum (HI-FBS, Thermo Fisher Scientific, Waltham, MA), 1% penicillin-streptomycin (PenStrep, 100X, FUJIFILM Wako) and 0.1% 2-mercaptoethanol (2-ME, 1000X, FUJIFILM Wako)].

A mouse T cell hybridoma with NFAT-GFP reporter gene, lacking endogenous TCR expression^27^, referred to as CT237, was a kind gift from the Yamasaki lab. CT237 cells were maintained in complete RPMI media [cRPMI; RPMI-1640 (FUJIFILM Wako) supplemented by 10% HI-FBS (Thermo Fisher Scientific, Waltham, MA), 1% PenStrep (100X, FUJIFILM Wako) and 0.1% 2-ME (1000X, FUJIFILM Wako)]. The mouse B cell line LK35.2 (ATCC-HB98), which expresses I-A^k^ MHC II, was used as the antigen presenting cell (APC) and was maintained in cDMEM. All cells were maintained in a humidified incubator with a 5% CO_2_ at 37 °C.

### Cloning

Wild-type and mutant 3A9 TCRɑ and β genes (nucleotide sequences are listed in table S1), synthesized by Integrated Device Technology (IDT), were inserted into retroviral vectors pMX-IRES-rat CD2 and pMX-IRES-human CD8, respectively, using In-Fusion snap assembly master mix kit (TAKARA Bio., Shiga, Japan, #638947) following the manufacturer’s protocol. The constructs were expanded, purified and validated by sequencing. Constructs were stored at -20 °C. Both retroviral vectors were a kind gift from the Yamasaki lab.

### Transfection/infection

Phoenix cells were plated in 6-well plates at a density of 3.5 × 10^5^ cells/well in 2 ml cDMEM. The following day, cells were transfected with TCRɑ and β plasmids in the same reaction, using PEI MAX (Polysciences) and incubated at 37 °C and 5% CO2 for 48 h. Retrovirus-containing supernatant was collected, filtered through a 0.22-µm filter, and concentrated by centrifugation. CT237 cells cultured in RPMI-1640 media, supplemented by 10% HI-FBS and 0.1% 2-ME, were then infected by the concentrated retrovirus supernatant along with DOTAP (Roche, #11202375001). Infected cells were left for expansion for 3-5 days before being purified based on the presence of CD3 using magnetic sorting.

After expansion cells were stained with APC/Cyanine7 anti-mouse CD3 antibodies (BioLegend, clone:17A2, #100221) and purified by positive selection from the cell suspension, using APC-conjugated MACS sorting magnetic beads (Miltenyi Biotec). Purification efficiency was assessed by flow cytometry and clones with >90% purity were used for further experiments.

### T cell stimulation

A total of 1 × 10^6^ purified 3A9 T cells and 1 × 10^6^ APCs were co-cultured in 96 well plates with a serial dilution of HEL peptide ranging from 0.016 to 1 μg/ml. Anti-mouse CD3 antibody (clone: 2C11, produced in-lab) was used to coat wells as a positive control. Plates were then incubated for different time points (2 h, 4 h, and 24 h) at 37 °C in a humidified incubator with 5% CO_2_. Each cell group was run in duplicates or triplicates per independent experiment.

After each time point, cells were stained with APC anti-human CD8a antibody (BioLegend, clone: RPA-T8, #301049) and incubated for 30 minutes on ice. Cells were washed with HBSS (1X, Gibco) twice then stained with 200 ul propidium iodide (PI, BioLegend, #421301). GFP expression level was assessed by the Attune NxT flow cytometer (Invitrogen). Flow cytometry data was analyzed using FlowJo v10.8 software. The baseline fluorescence intensity of negative controls without HEL peptide was used to set the gate placement. Lymphocytes gates were selected based on forward and side scatter plots, dead cells were excluded using PI, cells positive for APC anti-human CD8 antibody were selected as T cell gates from which GFP-positive T cells were assessed.

### Statistical analysis

Statistical tests were performed using GraphPad Prism 9 software. Comparing GFP expression level among multiple groups were performed using two-way ANOVA. Results were considered statistically significant when p-values were less than 0.05.

## Notes

### Competing Interest Statement

The authors have declared no competing interest.

### Summary of Updates

Expanded the discussion and improved the method section

## Bibliography

1. van der Merwe, P. A. & Dushek, O. Mechanisms for T cell receptor triggering. Nat Rev Immunol 11, 47–55 (2011).

2. Xu, C. et al. Regulation of T cell receptor activation by dynamic membrane binding of the CD3ε cytoplasmic tyrosine-based motif. Cell 135, 702–713 (2008).

3. Aivazian, D. & Stern, L. J. Phosphorylation of T cell receptor zeta is regulated by a lipid dependent folding transition. Nat Struct Biol 7, 1023–1026 (2000).

4. Sokurenko, E. V., Vogel, V. & Thomas, W. E. Catch-bond mechanism of force-enhanced adhesion: counterintuitive, elusive, but. widespread. Cell Host Microbe 4, 314–323 (2008).

5. Zhu, C., Chen, W., Lou, J., Rittase, W. & Li, K. Mechanosensing through immunoreceptors. Nature Publishing Group 1–10 (2019).

6. Feng, Y., Reinherz, E. L. & Lang, M. J. αβ T Cell Receptor Mechanosensing Forces out Serial Engagement. Trends Immunol 39, 596–609 (2018).

7. Hwang, W., Mallis, R. J., Lang, M. J. & Reinherz, E. L. The αβTCR mechanosensor exploits dynamic ectodomain allostery to optimize its ligand recognition site. Proc Natl Acad Sci U S A 117, 21336–21345 (2020).

8. Das, D. K. et al. Force-dependent transition in the T-cell receptor β-subunit allosterically regulates peptide discrimination and pMHC bond lifetime. Proc Natl Acad Sci U S A 112, 1517–1522 (2015).

9. Kim, S. T. et al. The alphabeta T cell receptor is an anisotropic mechanosensor. J Biol Chem 284, 31028–31037 (2009).

10. Lo, W. L. et al. Slow phosphorylation of a tyrosine residue in LAT optimizes T cell ligand discrimination. Nat Immunol 20, 1481–1493 (2019).

11. Al-Aghbar, M. A., Jainarayanan, A. K., Dustin, M. L. & Roffler, S. R. The interplay between membrane topology and mechanical forces in regulating T cell receptor activity. Commun Biol 5, 40 (2022).

12. Rangarajan, S. et al. Peptide-MHC (pMHC) binding to a human antiviral T cell receptor induces long-range allosteric communication between pMHC- and CD3-binding sites. The Journal of biological chemistry 293, 15991–16005 (2018).

13. Brazin, K. N. et al. The T Cell Antigen Receptor α Transmembrane Domain Coordinates Triggering through Regulation of Bilayer Immersion and CD3 Subunit Associations. Immunity 49, 829–841.e6 (2018).

14. Lanz, A. L. et al. Allosteric activation of T cell antigen receptor signaling by quaternary structure relaxation. Cell Rep 36, 109375 (2021).

15. Dong, D. et al. Structural basis of assembly of the human T cell receptor–CD3 complex. Nature 1–21 (2019).

16. Prakaash, D., Cook, G. P., Acuto, O. & Kalli, A. C. Multi-scale simulations of the T cell receptor reveal its lipid interactions, dynamics and the arrangement of its cytoplasmic region. PLoS Comput Biol 17, e1009232 (2021).

17. Pandey, P. R., Rozycki, B., Lipowsky, R. & Weikl, T. R. Structural variability and concerted motions of the T cell receptor-CD3 complex. eLife 10, (2021).

18. Sušac, L. et al. Structure of a fully assembled tumor-specific T cell receptor ligated by pMHC. Cell 185, 3201–3213.e19 (2022).

19. Newell, E. W. et al. Structural Basis of Specificity and Cross-Reactivity in T Cell Receptors Specific for Cytochrome c–I-Ek. J Immunol 186, 5823–5832 (2011).

20. He, Y. et al. Peptide–MHC Binding Reveals Conserved Allosteric Sites in MHC Class I- and Class II-Restricted T Cell Receptors (TCRs). Journal of Molecular Biology 432, 166697 (2020).

21. Mariuzza, R. A., Agnihotri, P. & Orban, J. The structural basis of T-cell receptor (TCR) activation: An enduring enigma. Journal of Biological Chemistry 295, 914–925 (2020).

22. Natarajan, K. et al. An allosteric site in the T-cell receptor Cβ domain plays a critical signalling role. Nature communications 8, 1–14 (2017).

23. Sasada, T. et al. Involvement of the TCR Cβ FG loop in thymic selection and T cell function. The Journal of experimental medicine 195, 1419–1431 (2002).

24. Touma, M. et al. The TCR C beta FG loop regulates alpha beta T cell development. J Immunol 176, 6812–6823 (2006).

25. Huang, J. et al. CHARMM36m: an improved force field for folded and intrinsically disordered proteins. Nature Publishing Group 14, 71–73 (2016).

26. Ho, W. Y., Cooke, M. P., Goodnow, C. C. & Davis, M. M. Resting and anergic B cells are defective in CD28-dependent costimulation of naive CD4+ T cells. The Journal of experimental medicine 179, 1539–1549 (1994).

27. Matsumoto, Y. et al. A TCR-like antibody against a proinsulin-containing fusion peptide ameliorates type 1 diabetes in NOD mice. Biochemical and Biophysical Research Communications 534, 680–686 (2021).

28. Chen, Y. et al. Cholesterol inhibits TCR signaling by directly restricting TCR-CD3 core tunnel motility. Mol Cell S1097–2765(22)00155 (2022).

29. Li, Q. et al. Polyunsaturated eicosapentaenoic acid changes lipid composition in lipid rafts. Eur J Nutr 45, 144–151 (2006).

30. Werlen, G. & Palmer, E. The T-cell receptor signalosome: a dynamic structure with expanding complexity. Curr Opin Immunol 14, 299–305 (2002).

31. Montixi, C. et al. Engagement of T cell receptor triggers its recruitment to low-density detergent-insoluble membrane domains. EMBO J 17, 5334–5348 (1998).

32. Xavier, R., Brennan, T., Li, Q., McCormack, C. & Seed, B. Membrane compartmentation is required for efficient T cell activation. Immunity 8, 723–732 (1998).

33. Wang, F., Beck-García, K., Zorzin, C., Schamel, W. W. A. & Davis, M. M. Inhibition of T cell receptor signaling by cholesterol sulfate, a naturally occurring derivative of membrane cholesterol. Nature immunology 17, 844–850 (2016).

34. Swamy, M. et al. A Cholesterol-Based Allostery Model of T Cell Receptor Phosphorylation. Immunity 44, 1091–1101 (2016).

35. Godoy-Hernandez, A. et al. Rapid and Highly Stable Membrane Reconstitution by LAiR Enables the Study of Physiological Integral Membrane Protein Functions. ACS Central Science (2023).

36. Banik, D. et al. Single Molecule Force Spectroscopy Reveals Distinctions in Key Biophysical Parameters of αβ T-Cell Receptors Compared with Chimeric Antigen Receptors Directed at the Same Ligand. J Phys Chem Lett 12, 7566–7573 (2021).

37. Das, D. K. et al. Pre-T Cell Receptors (Pre-TCRs) Leverage Vβ Complementarity Determining Regions (CDRs) and Hydrophobic Patch in Mechanosensing Thymic Self-ligands. The Journal of biological chemistry 291, 25292–25305 (2016).

38. Natarajan, A. et al. Structural Model of the Extracellular Assembly of the TCR-CD3 Complex. CellReports 14, 2833–2845 (2016).

39. Beddoe, T. et al. Antigen ligation triggers a conformational change within the constant domain of the αβ T cell receptor. Immunity 30, 777–788 (2009).

40. Hawse, W. F. et al. Cutting edge: Evidence for a dynamically driven T cell signaling mechanism. J Immunol 188, 5819–5823 (2012).

41. Brameshuber, M. et al. Monomeric TCRs drive T cell antigen recognition. Nat Immunol 19, 487–496 (2018).

42. Feng, Y. et al. Mechanosensing drives acuity of alphabeta T-cell recognition. Proc Natl Acad Sci U S A 114, E8204–E8213 (2017).

43. Friedl, P., den Boer, A. T. & Gunzer, M. Tuning immune responses: diversity and adaptation of the immunological synapse. Nature reviews. Immunology 5, 532–545 (2005).

44. Hopfield, J. J. Kinetic proofreading: a new mechanism for reducing errors in biosynthetic processes requiring high specificity. Proc Natl Acad Sci U S A 71, 4135–4139 (1974).

45. McKeithan, T. W. Kinetic proofreading in T-cell receptor signal transduction. Proc Natl Acad Sci U S A 92, 5042–5046 (1995).

46. Chatenoud, L. CD3-specific antibody-induced active tolerance: from bench to bedside. Nat Rev Immunol 3, 123–132 (2003).

47. Gaglia, J. & Kissler, S. Anti-CD3 Antibody for the Prevention of Type 1 Diabetes: A Story of Perseverance. Biochemistry 58, 4107–4111 (2019).

48. Pillarisetti, K. et al. A T-cell-redirecting bispecific G-protein-coupled receptor class 5 member D x CD3 antibody to treat multiple myeloma. Blood 135, 1232–1243 (2020).

49. Zimmermann, L. et al. A completely reimplemented MPI bioinformatics toolkit with a new HHpred server at its core. Journal of molecular biology 430, 2237–2243 (2018).

50. Schrödinger, L. L. C. The PyMOL Molecular Graphics System, Version 1.7.0.0. (2010).

51. Fiser, A. & Sali, A. ModLoop: automated modeling of loops in protein structures. Bioinformatics 19, 2500–2501 (2003).

52. Lis, M. et al. Bridging the gap between single-template and fragment based protein structure modeling using Spanner. Immunome Research 7, 1 (2011).

53. Jo, S., Kim, T., Iyer, V. G. & Im, W. CHARMM-GUI: a web-based graphical user interface for CHARMM. Journal of computational chemistry 29, 1859–1865 (2008).

54. Souza, P. C. T. et al. Martini 3: a general purpose force field for coarse-grained molecular dynamics. Nat Methods 1–7 (2021).

55. Thallmair, S., Vainikka, P. A. & Marrink, S. J. Lipid Fingerprints and Cofactor Dynamics of Light-Harvesting Complex II in Different Membranes. Biophysical journal 116, 1446–1455 (2019).

56. Poma, A. B., Cieplak, M. & Theodorakis, P. E. Combining the MARTINI and Structure-Based Coarse-Grained Approaches for the Molecular Dynamics Studies of Conformational Transitions in Proteins. Journal of Chemical Theory and Computation 13, 1366–1374 (2017).

57 . Kroon, P. C. Automate, aggregate, assemble. (2020).

58. Abraham, M. J. et al. GROMACS: High performance molecular simulations through multi-level parallelism from laptops to supercomputers. SoftwareX 1, 19–25 (2015).

59. de Jong, D. H., Baoukina, S., Ingólfsson, H. I. & Marrink, S. J. Martini straight: Boosting performance using a shorter cutoff and GPUs. Computer Physics Communications 199, 1–7 (2016).

60. Bussi, G., Donadio, D. & Parrinello, M. Canonical sampling through velocity rescaling. The Journal of Chemical Physics 126, 014101 (2007).

61. Berendsen, H. J. C., Postma, J. P. M., van Gunsteren, W. F., DiNola, A. & Haak, J. R. Molecular dynamics with coupling to an external bath. The Journal of Chemical Physics 81, 3684–3690 (1984).

62. Lee, J. et al. CHARMM-GUI input generator for NAMD, GROMACS, AMBER, OpenMM, and CHARMM/OpenMM simulations using the CHARMM36 additive force field. Journal of chemical theory and computation 12, 405-413 (2016).

63. Darden, T., York, D. & Pedersen, L. Particle mesh Ewald: An N⋅ log (N) method for Ewald sums in large systems. The Journal of Chemical Physics 98, 10089–10092 (1993).

64. Essmann, U. et al. A smooth particle mesh Ewald method. The Journal of Chemical Physics 103, 8577–8593 (1995).

65. Nosé, S. A unified formulation of the constant temperature molecular dynamics methods. The Journal of Chemical Physics 81, 511–519 (1984).

66. Hoover, W. G. Canonical dynamics: Equilibrium phase-space distributions. Physical review A 31, 1695 (1985).

67. Parrinello, M. & Rahman, A. Polymorphic transitions in single crystals: A new molecular dynamics method. Journal of Applied Physics 52, 7182–7190 (1981).

68. Wassenaar, T. A., Ingólfsson, H. I., Böckmann, R. A., Tieleman, D. P. & Marrink, S. J. Computational Lipidomics with insane: A Versatile Tool for Generating Custom Membranes for Molecular Simulations. Journal of Chemical Theory and Computation 11, 2144–2155 (2015).

69. Gowers, R. J. et al. MDAnalysis: A Python package for the rapid analysis of molecular dynamics simulations. Proceedings of the 15th Python in Science Conference 102-109 (2016).

70. Michaud-Agrawal, N., Denning, E. J., Woolf, T. B. & Beckstein, O. MDAnalysis: a toolkit for the analysis of molecular dynamics simulations. Journal of Computational Chemistry 32, 2319–2327 (2011).

71. Humphrey, W., Dalke, A. & Schulten, K. VMD: Visual molecular dynamics. Journal of Molecular Graphics 14, 33–38 (1996).

72. Goddard, T. D. et al. UCSF ChimeraX: Meeting modern challenges in visualization and analysis. Protein Sci 27, 14–25 (2018).

73. Pettersen, E. F. et al. UCSF ChimeraX: Structure visualization for researchers, educators, and developers. Protein Sci 30, 70–82 (2021).

