## Supplementary material for "TCR-pMHC complex formation triggers CD3 dynamics": Suppl Figure S2

### Supplemental Figure S2: Contact plots

The barplots in this document describe the interactions summed over all simulations between individual CD3 proteins and the TCRα and TCRβ chains. Contacts in the TCR-CD3 simulations are shown in a green and contacts from the TCR-CD3-pMHC simulations in purple. Within each bar there is a color gradient, in which each shade represents a single simulation. Residue numbers of respecitively the TCRα and TCRβ chains are on the x-axis. The y-axis shows the culmulative number of interactions. Note that the y-axis varies between plots.

Panel A: CD3$\zeta^{\varepsilon}$-TCRα
Panel B: CD3$\zeta^{\varepsilon}$-TCRβ
Panel C: CD3$\zeta^{\gamma}$-TCRα
Panel D: CD3$\zeta^{\gamma}$-TCRβ
Panel E: CD3δ-TCRα
Panel F: CD3δ-TCRβ
Panel G: CD3$\varepsilon^{\delta}$-TCRα
Panel H: CD3$\varepsilon^{\delta}$-TCRβ
Panel I: CD3$\varepsilon^{\gamma}$-TCRα
Panel J: CD3$\varepsilon^{\gamma}$-TCRβ
Panel K: CD3γ-TCRα
Panel L: CD3γ-TCRβ

A

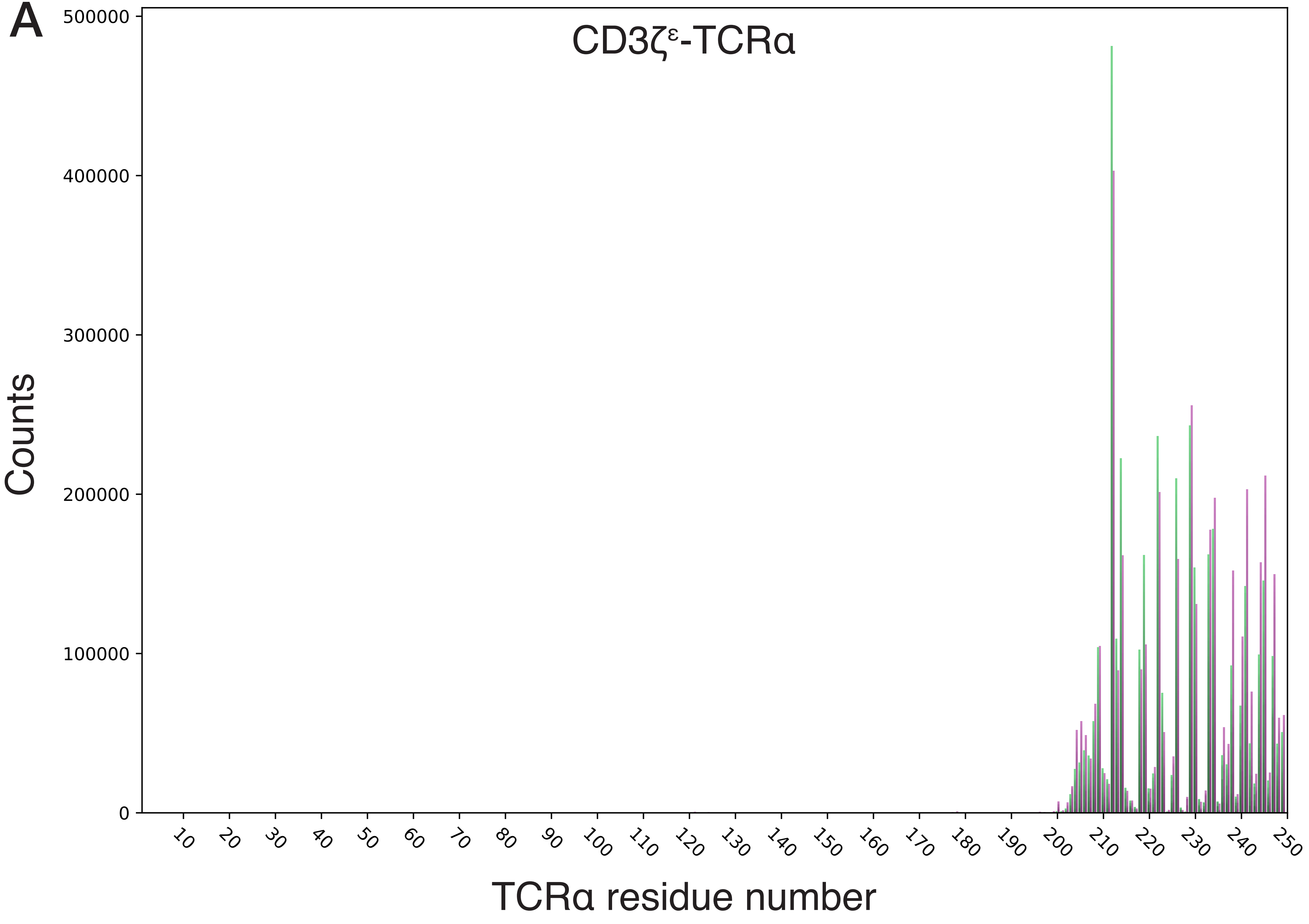
CD3ζ^ε-TCRα
Counts
TCRα residue number
0
100000
200000
300000
400000
500000
10
20
30
40
50
60
70
80
90
100
110
120
130
140
150
160
170
180
190
200
210
220
230
240
250

B

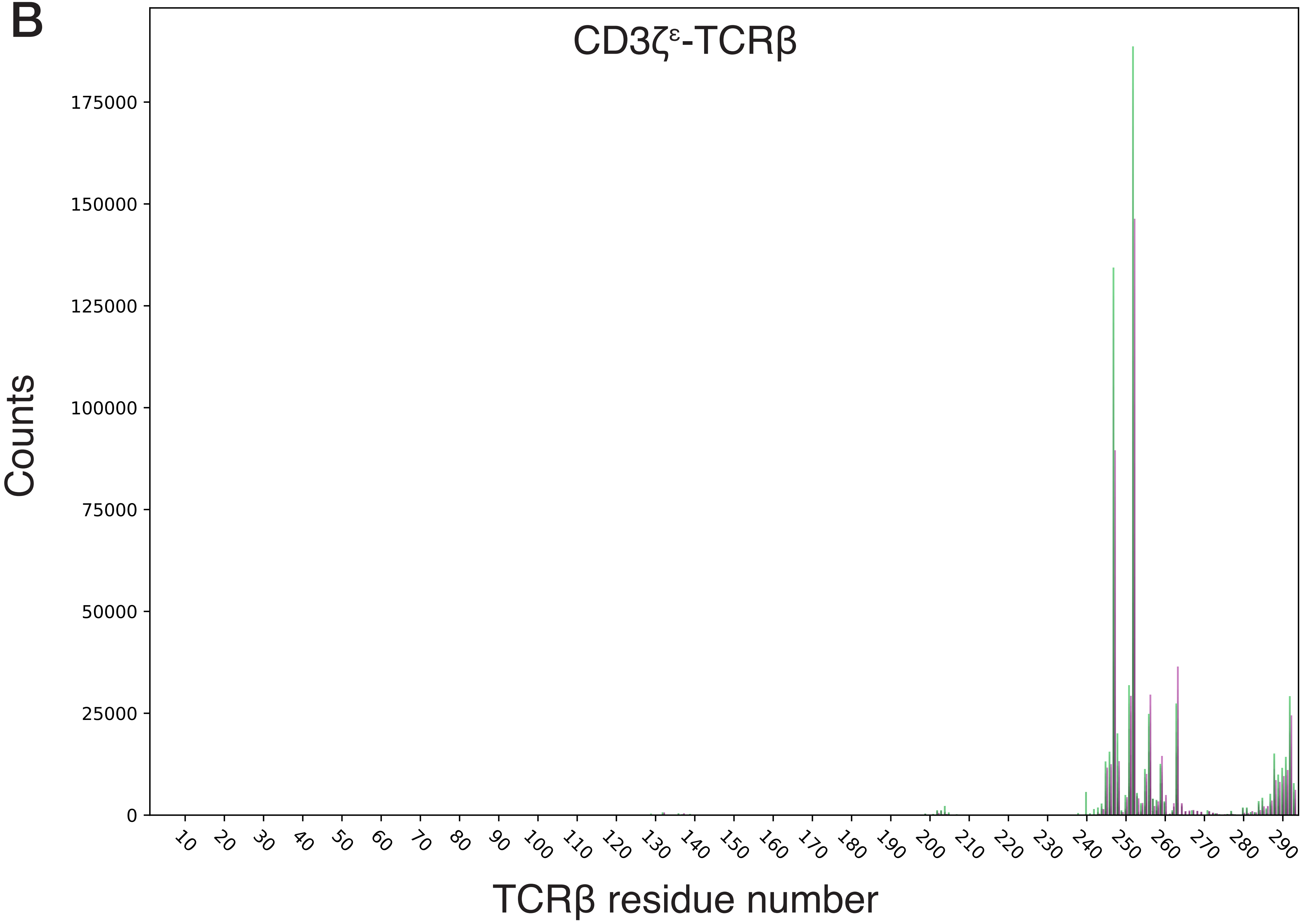
CD3ζᵋ-TCRβ
Counts
175000
150000
125000
100000
75000
50000
25000
0
10
20
30
40
50
60
70
80
90
100
110
120
130
140
150
160
170
180
190
200
210
220
230
240
250
260
270
280
290
TCRβ residue number

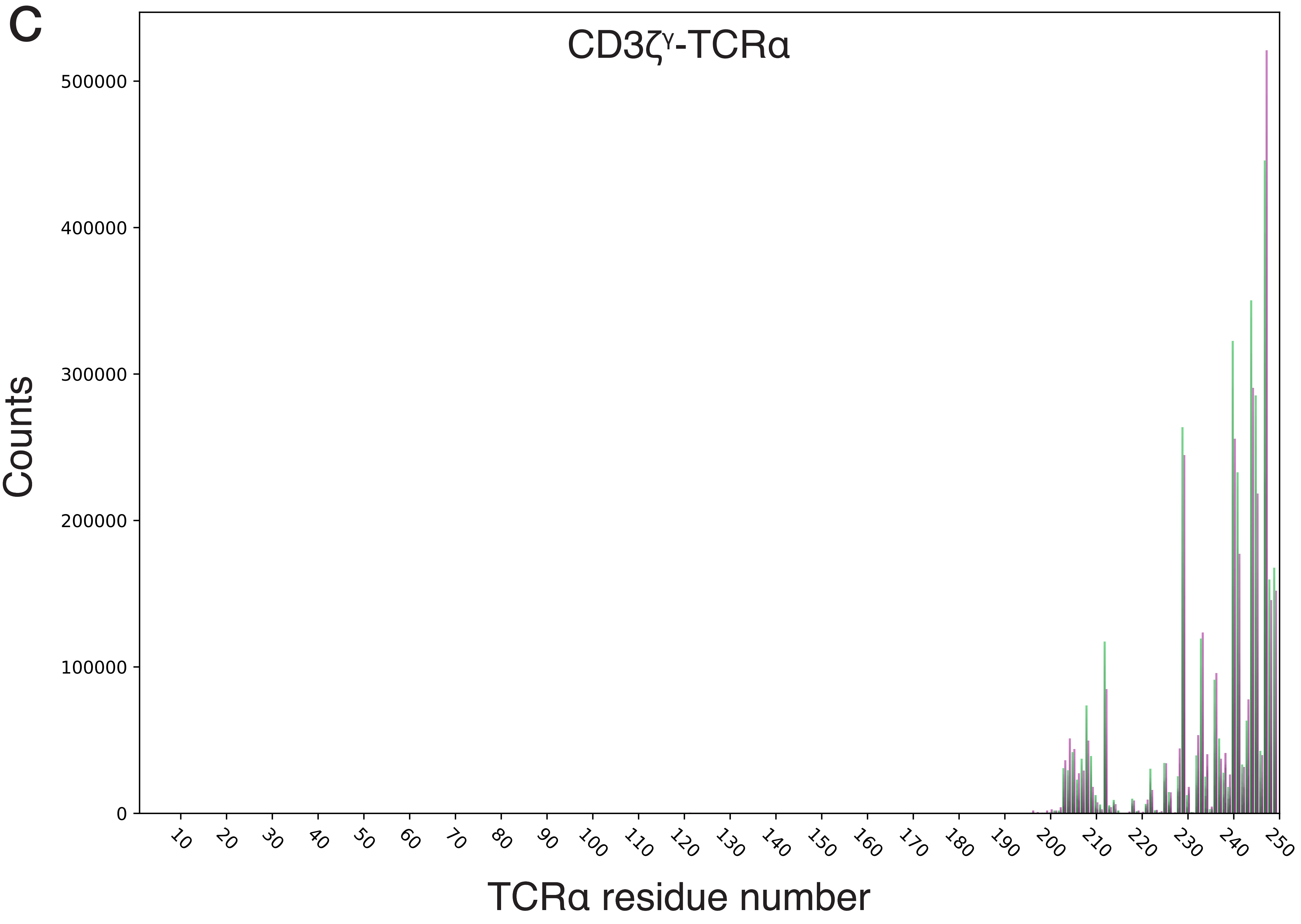
C
CD3ζγ-TCRα
Counts
500000
400000
300000
200000
100000
0
10
20
30
40
50
60
70
80
90
100
110
120
130
140
150
160
170
180
190
200
210
220
230
240
250
TCRα residue number

D

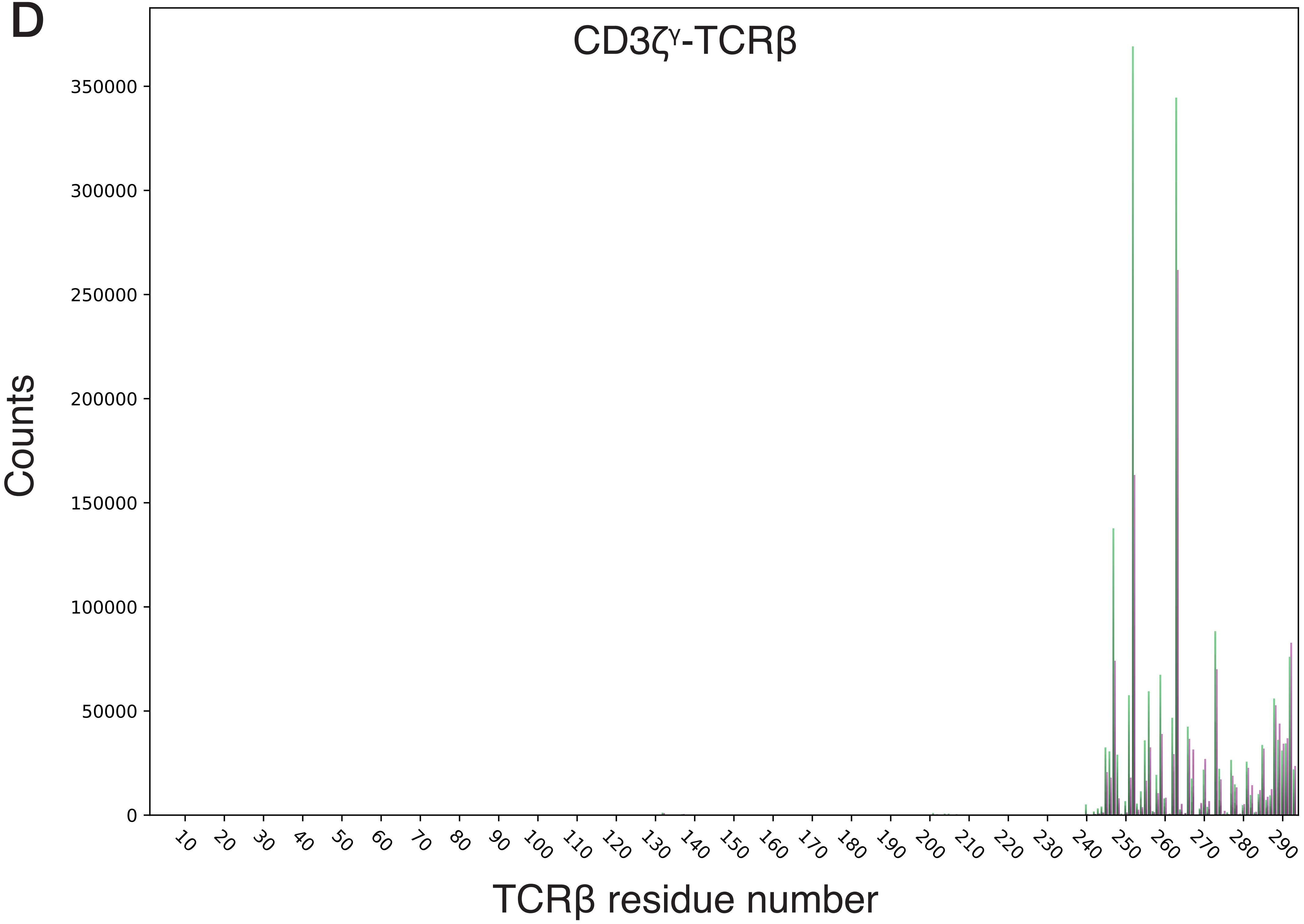

CD3ζγ-TCRβ
Counts
350000
300000
250000
200000
150000
100000
50000
0
10
20
30
40
50
60
70
80
90
100
110
120
130
140
150
160
170
180
190
200
210
220
230
240
250
260
270
280
290
TCRβ residue number

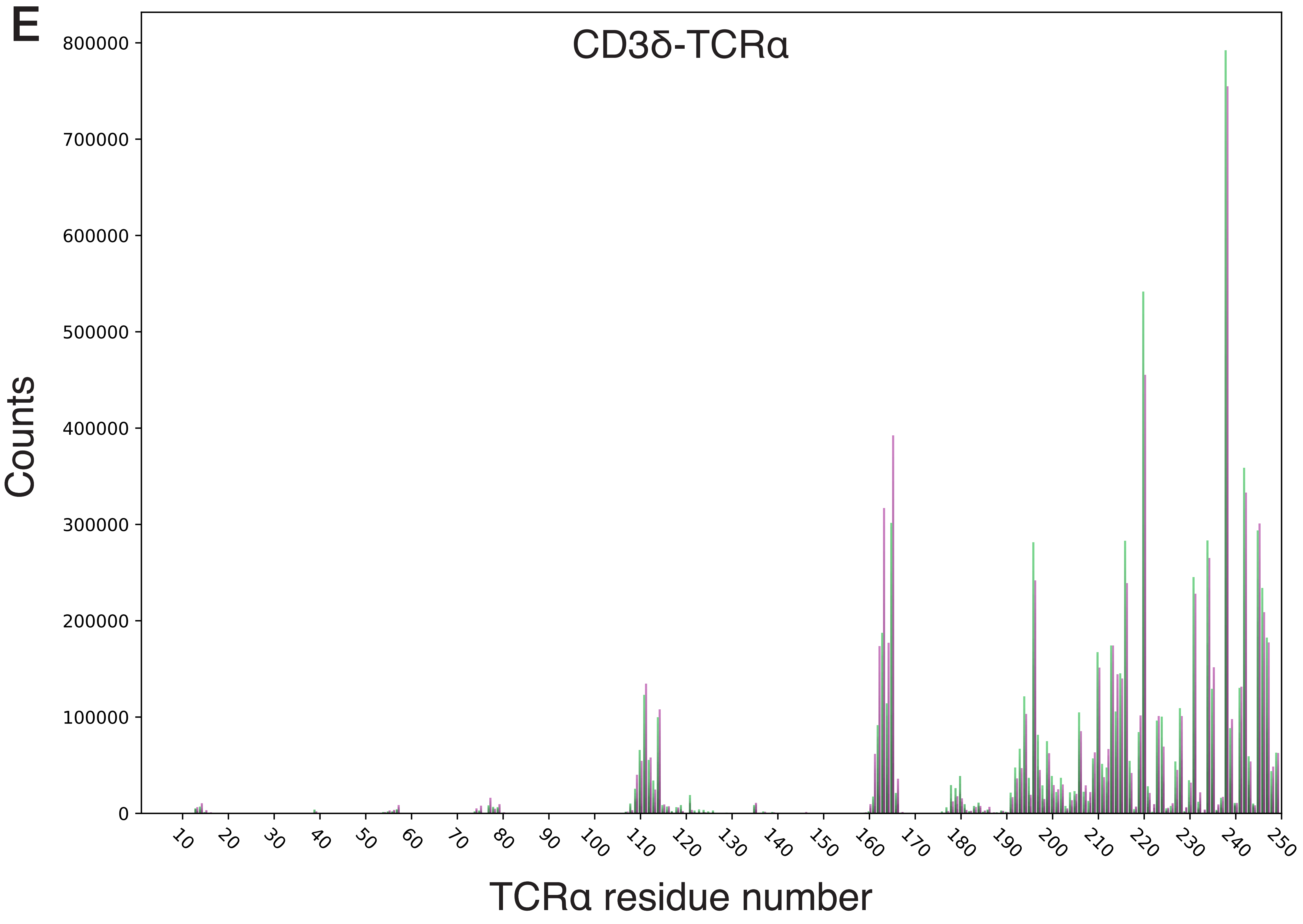
E
CD3δ-TCRα
Counts
800000
700000
600000
500000
400000
300000
200000
100000
0
10
20
30
40
50
60
70
80
90
100
110
120
130
140
150
160
170
180
190
200
210
220
230
240
250
TCRα residue number

F

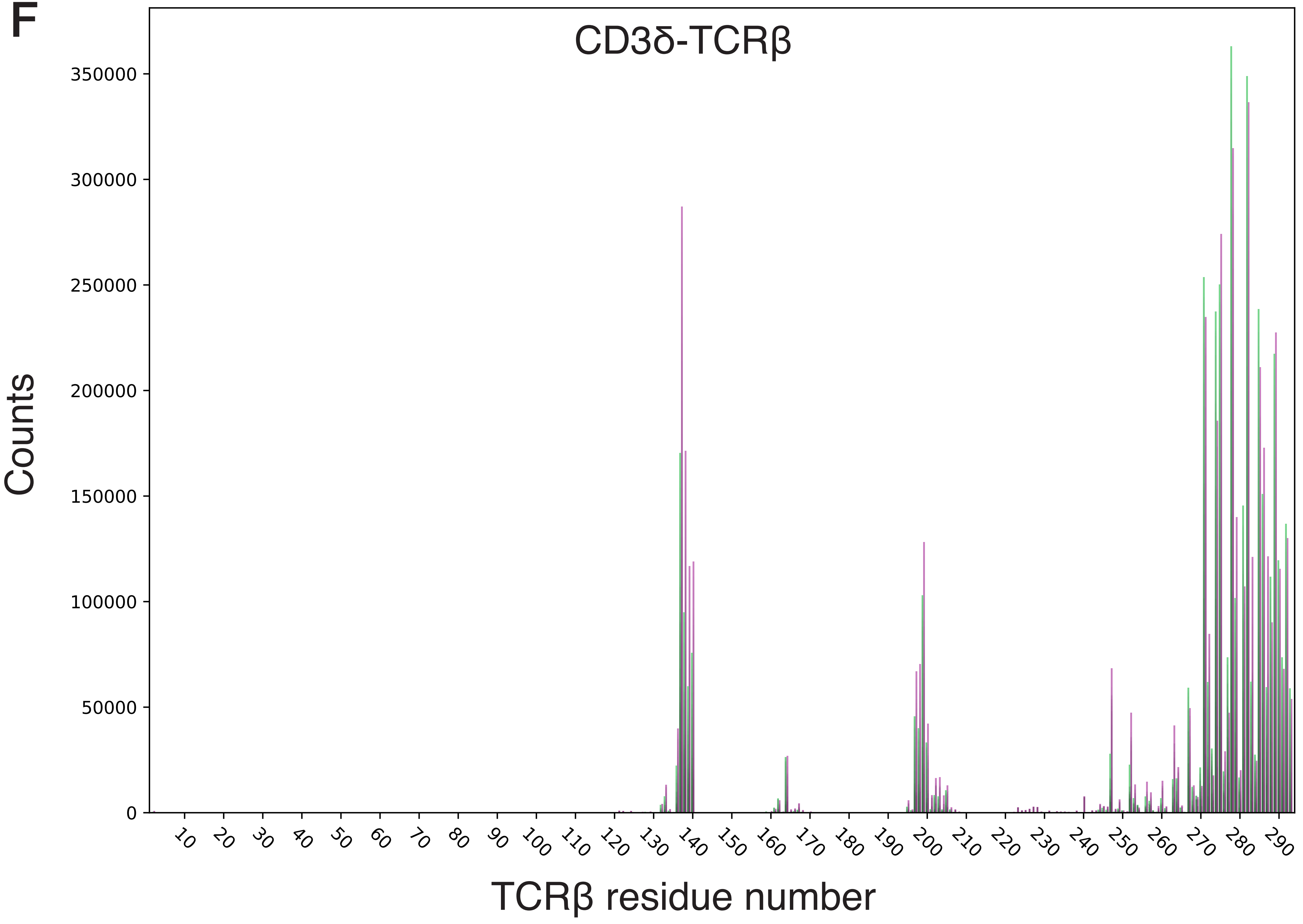

CD3δ-TCRβ
Counts
350000
300000
250000
200000
150000
100000
50000
0
10
20
30
40
50
60
70
80
90
100
110
120
130
140
150
160
170
180
190
200
210
220
230
240
250
260
270
280
290
TCRβ residue number

G

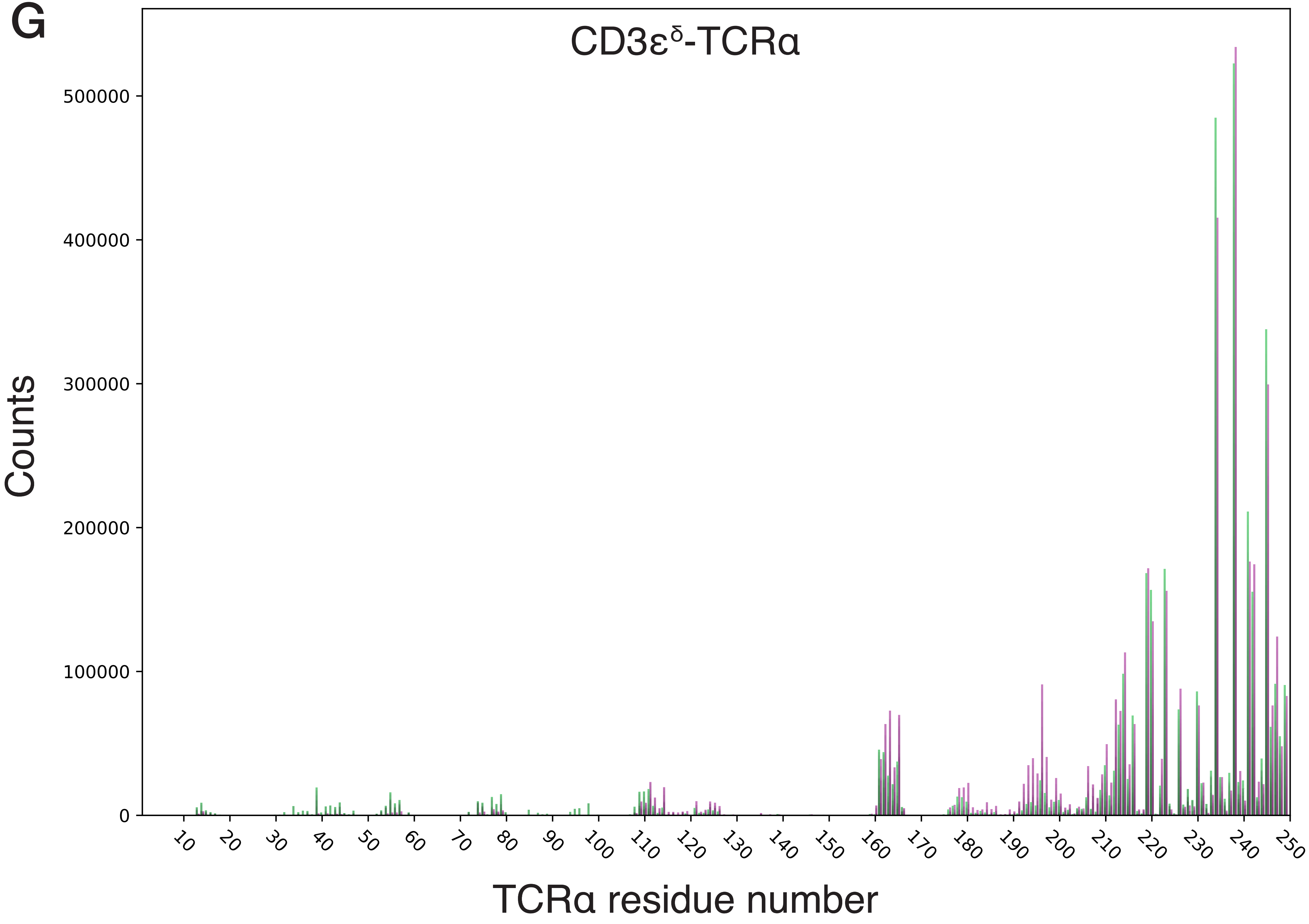
CD3ε^δ-TCRα
Counts
0
100000
200000
300000
400000
500000
10
20
30
40
50
60
70
80
90
100
110
120
130
140
150
160
170
180
190
200
210
220
230
240
250
TCRα residue number

H

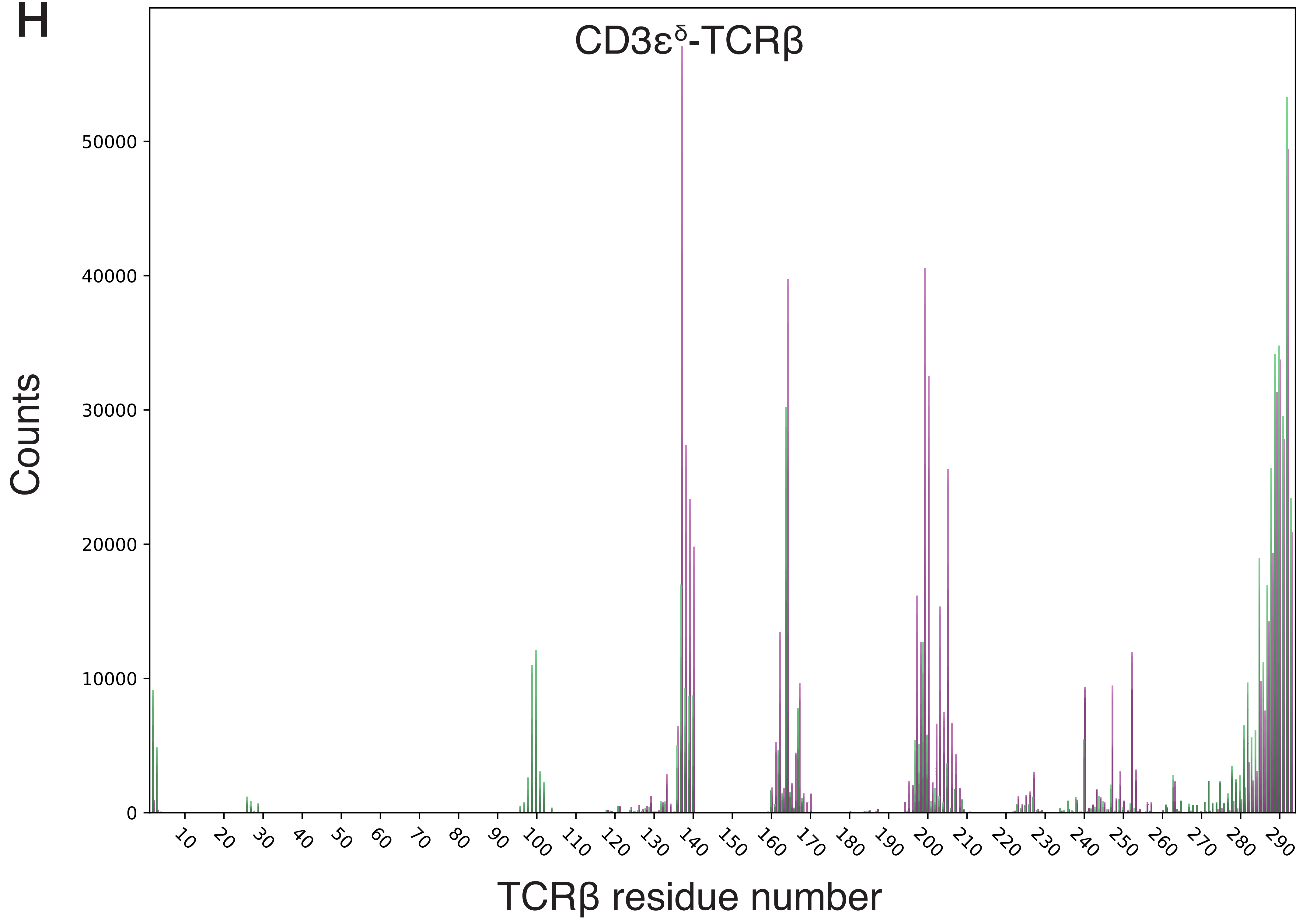

CD3ε$^{\delta}$-TCRβ
Counts
TCRβ residue number
0
10000
20000
30000
40000
50000
10 20 30 40 50 60 70 80 90 100 110 120 130 140 150 160 170 180 190 200 210 220 230 240 250 260 270 280 290

I

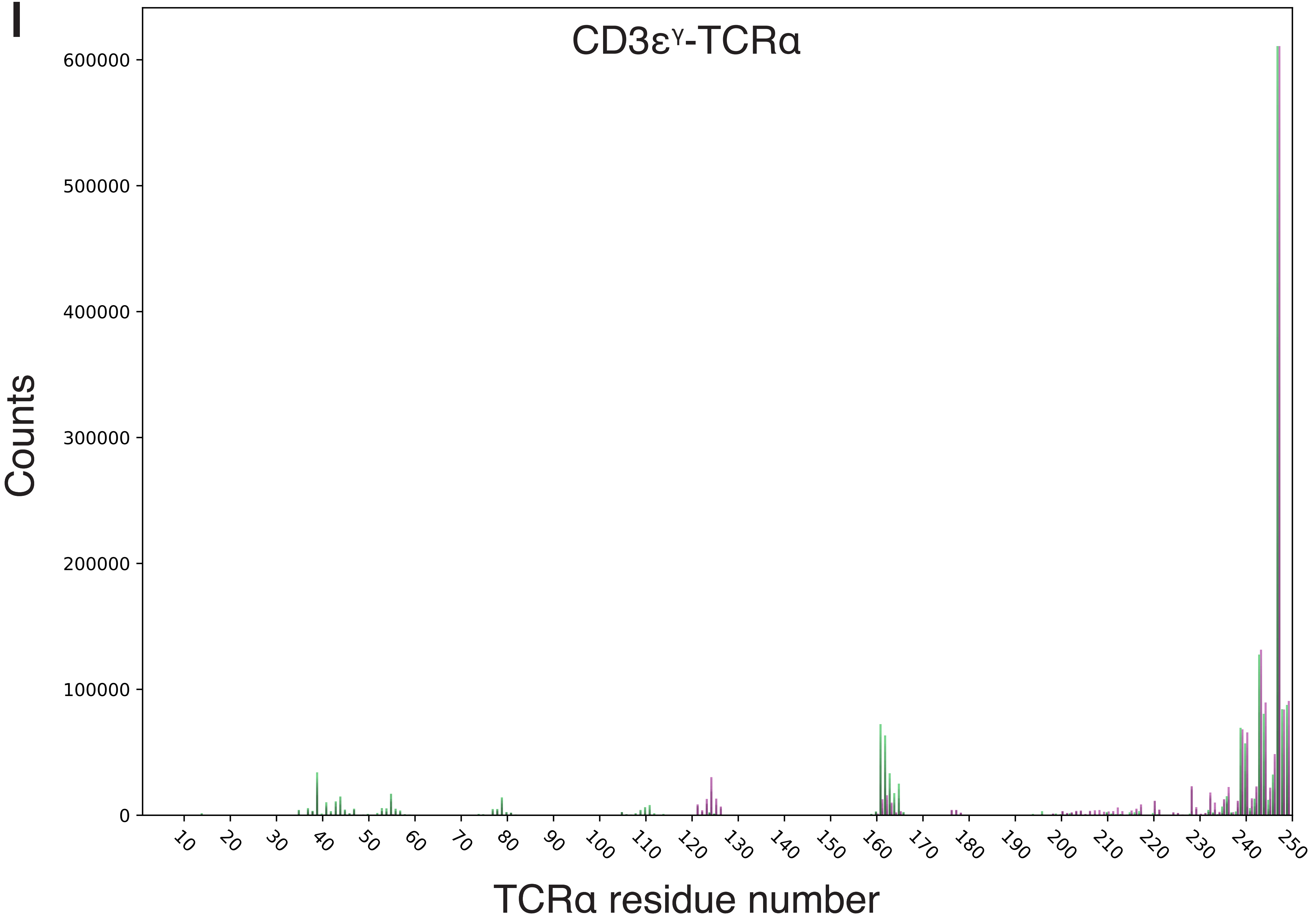

CD3εγ-TCRα
Counts
600000
500000
400000
300000
200000
100000
0
10
20
30
40
50
60
70
80
90
100
110
120
130
140
150
160
170
180
190
200
210
220
230
240
250
TCRα residue number

J

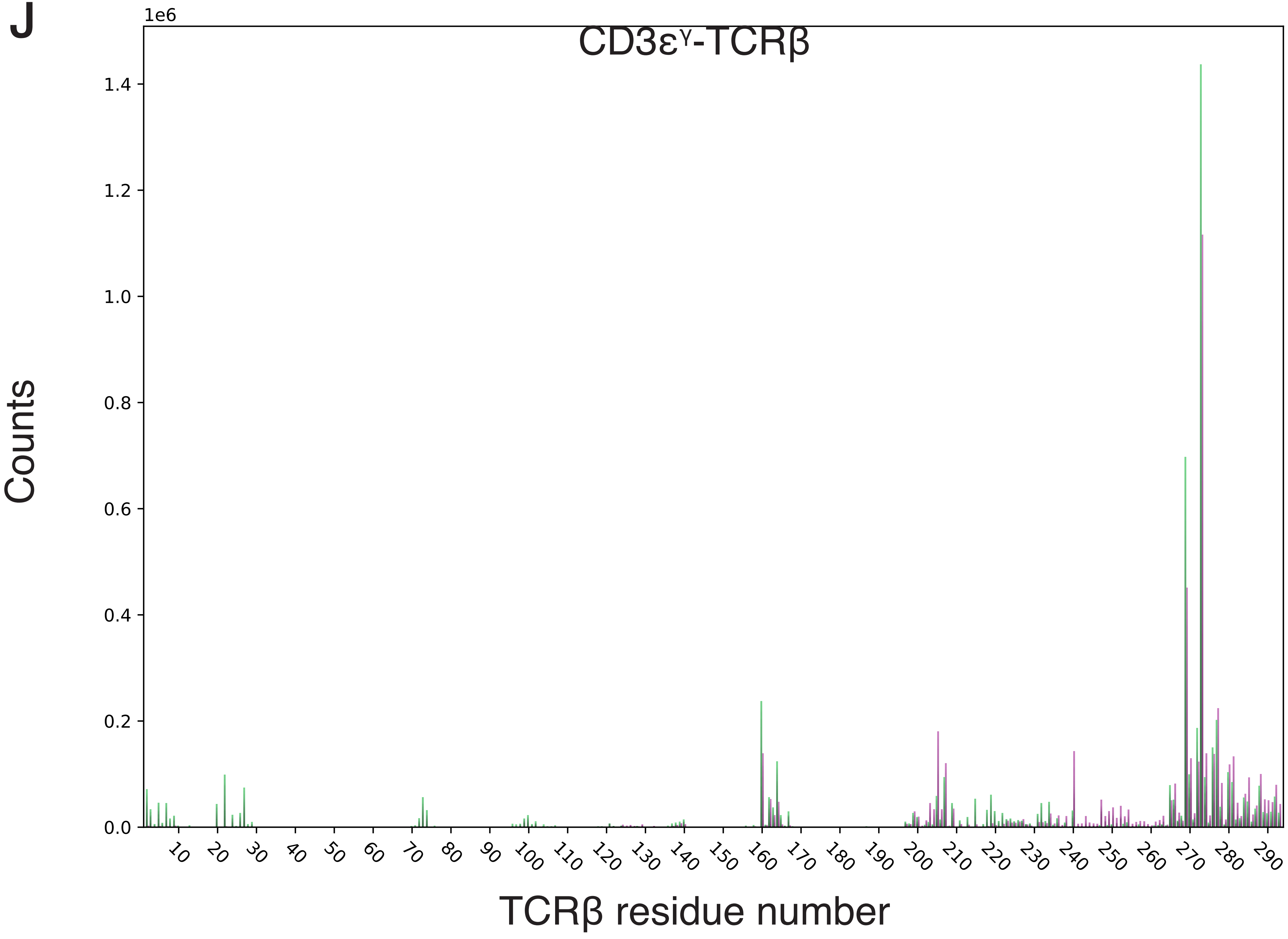
1e6
CD3εγ-TCRβ
Counts
1.4
1.2
1.0
0.8
0.6
0.4
0.2
0.0
10
20
30
40
50
60
70
80
90
100
110
120
130
140
150
160
170
180
190
200
210
220
230
240
250
260
270
280
290
TCRβ residue number

K

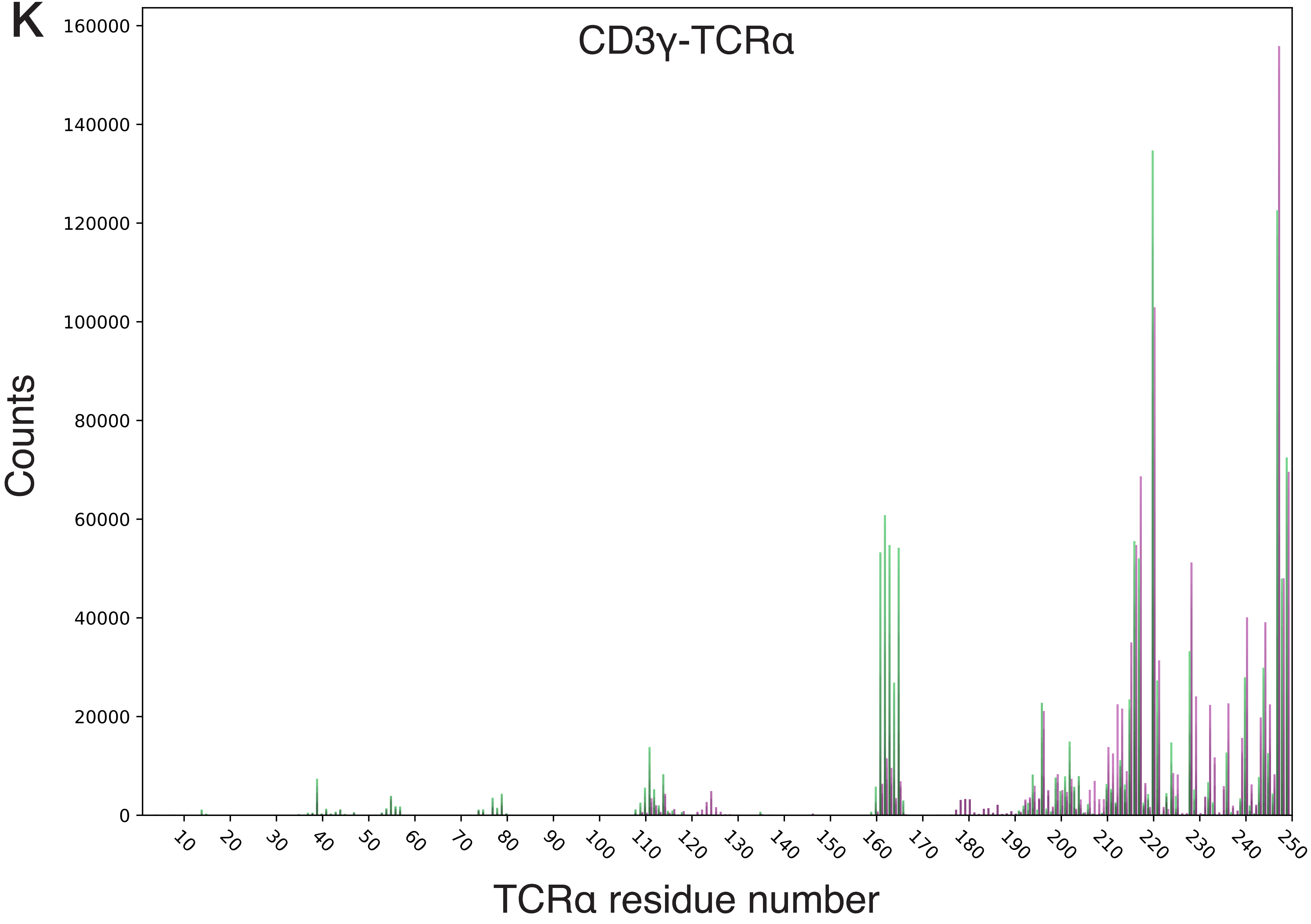
CD3γ-TCRα
Counts
160000
140000
120000
100000
80000
60000
40000
20000
0
10
20
30
40
50
60
70
80
90
100
110
120
130
140
150
160
170
180
190
200
210
220
230
240
250
TCRα residue number

L

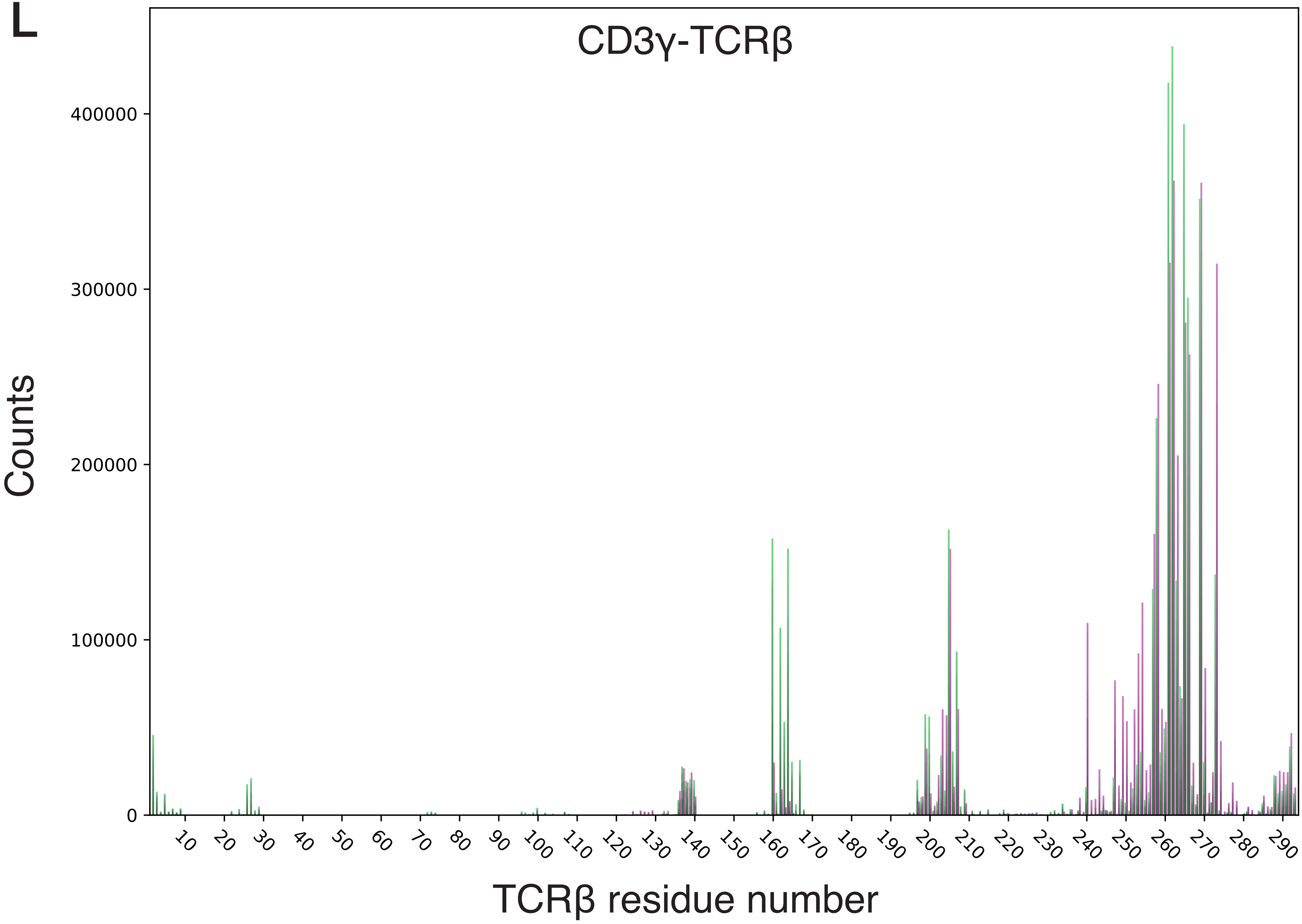
CD3γ-TCRβ
Counts
400000
300000
200000
100000
0
10 20 30 40 50 60 70 80 90 100 110 120 130 140 150 160 170 180 190 200 210 220 230 240 250 260 270 280 290
TCRβ residue number
