## Supplementary material for "TCR-pMHC complex formation triggers CD3 dynamics": Suppl Text & Suppl Figures S1,S3-S8

### Supplemental Text

*TCRα AB loop, the TCRβ H3 helix, the TCRβ FG loop and the CD3ζζ chains*

Long range perturbations of the TCR upon pMHC binding were previously observed in three regions: the TCRα AB loop, the TCRβ H3 helix, and the TCRβ FG loop[1–3]. In agreement with these experimental measurements, we also observed differences in these regions in our simulations upon pMHC binding. Both the TCRα AB loop and the TCRβ H3 helix had increased interactions with the CD3ε chains in the TCR-CD3-pMHC systems. In the TCR-CD3-pMHC simulations, the TCRα AB loop predominantly contacted CD3$\varepsilon^{\gamma}$ (Figure 6A), and the TCRβ H3 helix made more contacts with CD3$\varepsilon^{\delta}$ (Figure 6B). The TCRα AB loop and the TCRβ H3 helix were inaccessible to CD3$\varepsilon^{\gamma}$ and CD3$\varepsilon^{\delta}$ in the cryo-EM structure; this makes interactions between these regions unlikely in TCR-CD3 systems because CD3 movement is restricted. Upon pMHC binding, interactions between the TCR EC and the CD3 proteins were decoupled, which increased movement of the CD3ε chains around previously inaccessible areas of the TCR.

The TCRβ FG loop is important for TCR triggering and changes upon pMHC binding[3–5]. We observed that the TCRβ FG loop contacted CD3$\varepsilon^{\gamma}$ in the TCR-CD3 simulations more than in the TCR-CD3-pMHC simulations (Figure 6C). The TCRβ FG loop blocked CD3εγ diffusion around TCRβ; however, upon pMHC binding, the TCRβ FG loop extended upward, which allowed diffusion of CD3εγ. These observations suggest that the TCRβ FG loop prevents diffusion of the CD3εγ dimer when the TCR is unbound, thereby acting as a gatekeeper.

The TCRα chain lacks an equivalent TCRβ FG loop, which suggests that the CD3 dimer next to the TCRα chain – CD3εδ – may occasionally move alongside the TCRα chain; whereas, an equivalent movement by CD3εγ is blocked by the TCRβ FG loop. This explains the asymmetry observed in the CD3ε iso-occupancy map of the TCR-CD3 simulations (Figure 2A). The iso-occupancy maps in the TCR-CD3 simulation lack density specifically around CD3$\zeta^{\gamma}$, but there is significant density around CD3$\zeta^{\varepsilon}$ – the CD3ζ adjacent to CD3$\varepsilon^{\delta}$. This also can be observed in the contacts between the TCRα and the CD3ε chains (Figure S3), but the differences between the TCR-CD3 and TCR-CD3-pMHC simulations were much smaller compared to those between TCRβ and CD3εγ (Figure 3).

Lanz et al. observed a weakening in TCR-CD3ζ interactions upon TCR-pMHC binding[6]. We observe a similar effect in our simulations– a decrease in TCR-CD3ζ interactions upon TCR-pMHC binding (Figure S4). The decrease in contacts is especially clear in case of the TCRβ chain, although we cannot explain this asymmetry in the CD3ζζ contacts between the TCRα and TCRβ chains.

### Supplemental Figures S1, S3-S8

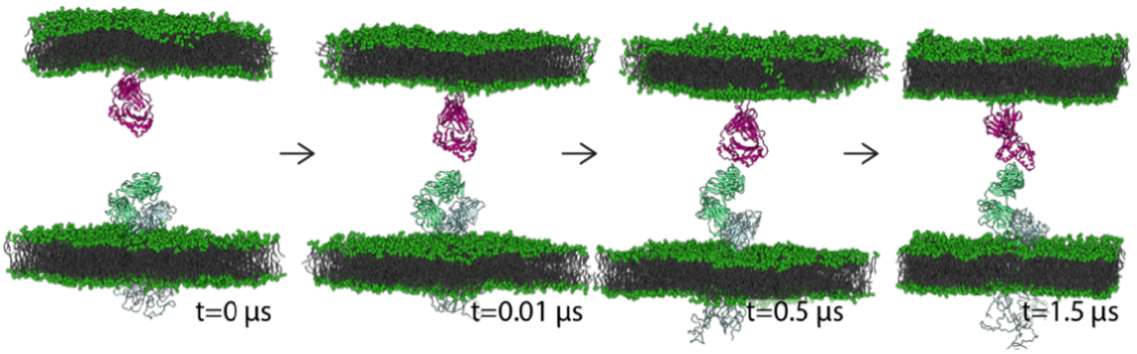


**Figure S1 Preparatory simulations of the TCR-CD3-pMHC systems.** pMHC was bound to TCR in a 1.5-µs preparatory simulation using a set of distance restraints based on the Newell structure[7]. A low force constant of 1 kJ mol$^{-1}$ nm$^{-2}$ was used to prevent deformations.

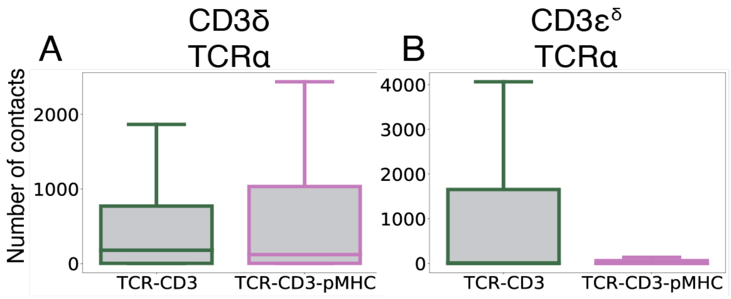


**Figure S3 TCR CD3 co-receptor contacts.** The number of contacts between the TCRα variable domain and the indicated CD3 chains in the TCR-CD3 (green box plots) and TCR-CD3-pMHC (purple box plots). Note that the y-axis varies between the two plots. For visibility, outliers are not shown. Please refer to Figure S2 for the contacts specified per residue.

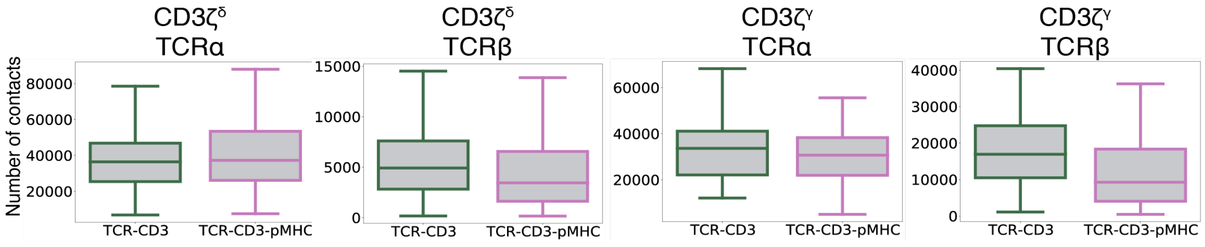


**Figure S4 TCR-CD3ζ chain contacts.** The number of contacts between the full TCRα or TCRβ chains and the indicated CD3ζ chains in the TCR-CD3 (green box plots) and TCR-CD3-pMHC (purple box plots). Note that the y-axis varies between the four plots. For visibility, outliers are not shown. Please refer to Figure S2 for the contacts specified per residue.

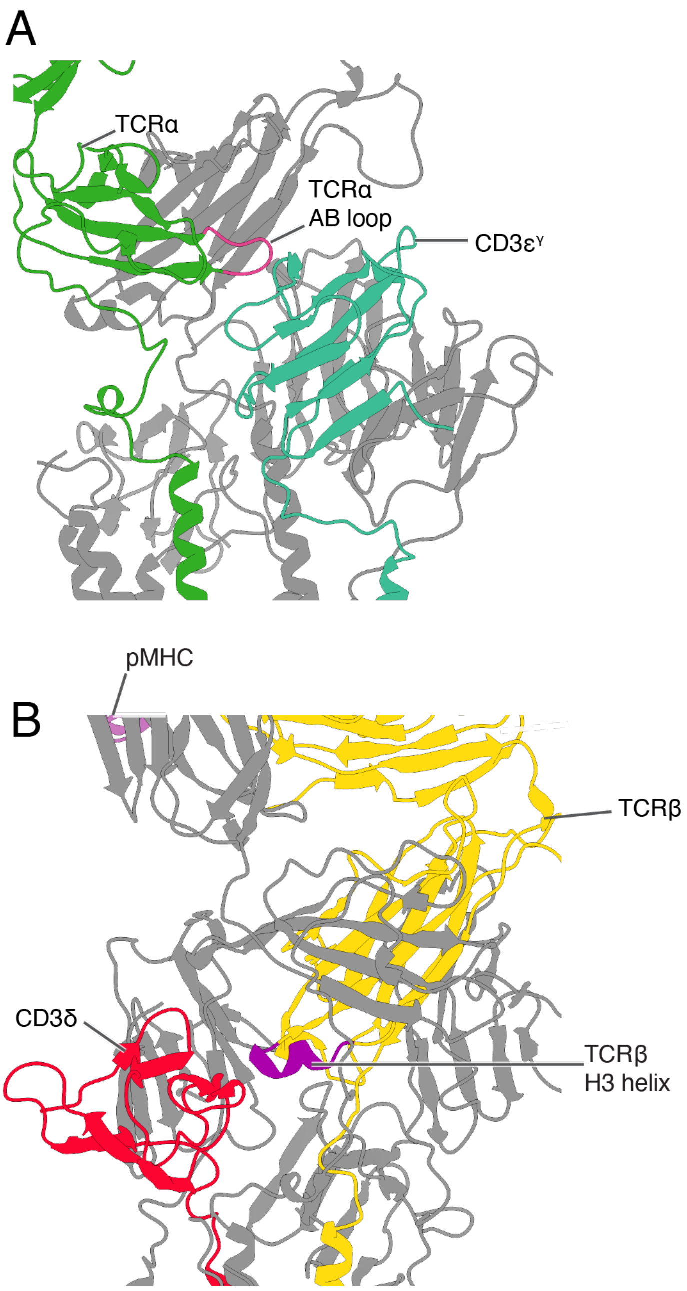


**Figure S5: CD3 proteins interact with the TCRα AB loop and TCRβ H3 helix upon TCR-pMHC binding.** Snapshots of the TCR-CD3-pMHC simulations illustrating how CD3$\varepsilon^{\gamma}$ interacts with the TCRα AB loop (panel A) and CD3δ with the TCRβ H3 helix (panel B) upon TCR-pMHC binding. In the TCR-CD3 simulations, the CD3 proteins cannot reach the TCRα AB loop or TCRβ H3 helix, but after pMHC binding, the CD3 proteins can diffuse around the TCR. TCRα is shown in green, TCRβ in yellow, the TCRα AB loop in fuchsia, the TCRβ H3 helix in purple, CD3$\varepsilon^{\gamma}$ in light green, CD3δ in red and the pMHC in light pink. Other protein chains are colored grey for clarity. The pMHC is not visible in panel A, and only barely visible in panel B.

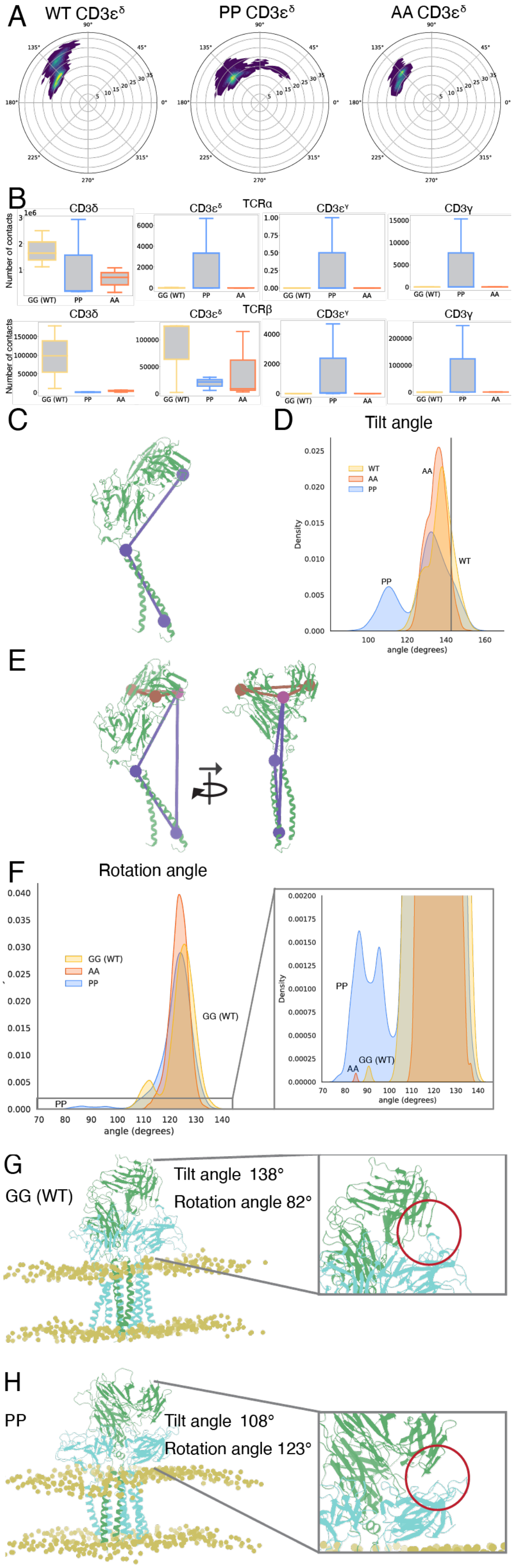

A
WT CD3εδ
PP CD3εδ
AA CD3εδ
B
CD3δ
CD3εδ
TCRα
CD3εγ
CD3γ
Number of contacts
GG (WT)
PP
AA
TCRβ
C
D
Tilt angle
WT
AA
PP
Density
angle (degrees)
E
F
Rotation angle
GG (WT)
AA
PP
angle (degrees)
Density
G
GG (WT)
Tilt angle 138°
Rotation angle 82°
H
PP
Tilt angle 108°
Rotation angle 123°

**Figure S6 Characterization of the atomistic TCR simulations.** (A) Polar plots of the CD3ε$^{\delta}$ movement around TCRβ in the atomistic TCR simulations of wild type (WT), double proline mutant (PP) and double alanine mutant (AA). (B) TCR-CD3 contacts in the atomistic simulations. The number of contacts between the TCRα (upper row) and TCRβ (bottom row) variable regions and the CD3δ, CD3ε$^{\delta}$, CD3ε$^{\gamma}$ and CD3γ chains. Wild-type boxplot in yellow, double proline mutant (PP) in blue and double alanine mutant in orange (AA). There is an increase in TCRα-CD3ε$^{\delta}$ interactions in the double proline mutant, but the total number of contacts between TCRαβ and the CD3δε is decreased in this mutant. Note that the axes have different scales. (C) Atoms used for the tilt angle calculation, which were TCRβ asparagine 97, TCRβ serine 249 and TCRβ valine 281. (D) Tilt of the TCR EC (TCRβ EC-TCRβ TM angle) in the atomistic simulations**.** Tilt angle distribution of the wild-type TCR (GG (WT), yellow), double proline mutant (PP, blue) and double alanine mutant (AA, orange). The angle of the TCR in the published cryo-EM structure[8] (vertical line) was 144°. (E) Illustration of the two planes that were used for the calculation of the rotation angle. Plane 1 (blue) was defined by the Cα atoms used for the tilt TCR EC tilt calculation: asparagine 97, serine 249 and valine 281 of the TCR β chain. Plane 2 (brown) was defined by the Cα atoms of TCRβ asparagine 97, TCRβ glycine 14 and TCRα serine 18. (F) Rotation of the TCR EC in the atomistic simulations. Distribution of the rotation angle of the TCR EC in wild type (GG (WT)), double proline mutant (PP) and double alanine mutant (AA) in the atomistic simulations with a zoom version of the plot (*right*). The proline distribution has a shoulder ~75-100°, which is virtually absent in wild type. Snapshots of the wild-type (G) and double proline simulations (H). Although wild type can restrain the CD3δ via contacts between TCRα and the CD3δ FG loop, the proline mutant is not able to, as shown in the red encircled area. Although the proline mutant adopts a smaller tilt angle (TCRβ EC with the TCRβ TM) of 108° (proline) vs. 138° (wild type) in this snapshot, it is due to the rotation of the TCR EC 82° (proline) vs. 123° (wild type) that the proline mutant is unable to retain the CD3 proteins. The TCR (green), CD3 proteins (cyan), and lipid headgroups (light brown) are shown

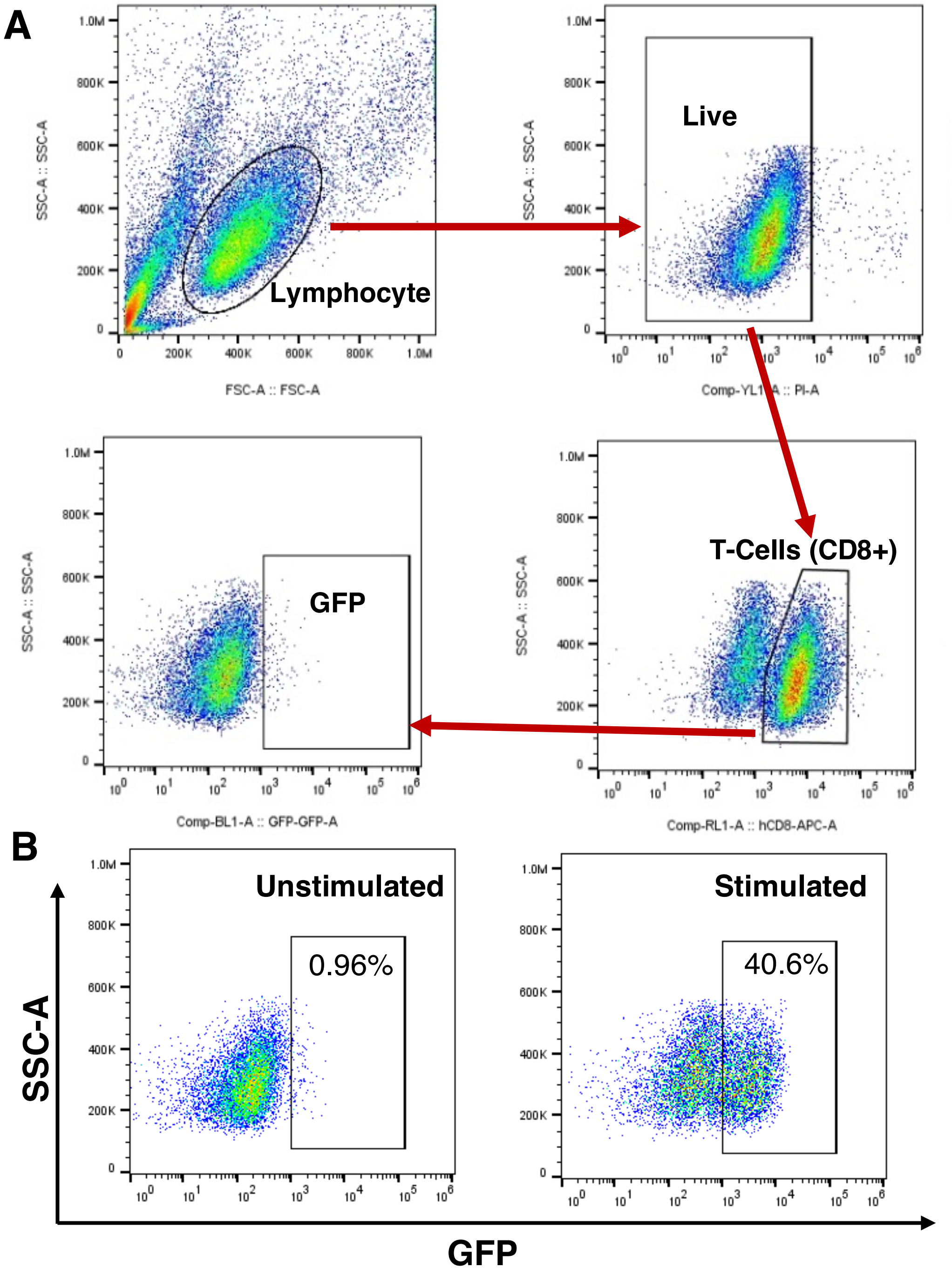


**Figure S7. Flow cytometry gating strategy for identifying GFP-positive T cells after co-culturing with APCs and HEL peptide. (A)** Flow cytometry gating strategy: the baseline fluorescence intensity of negative controls without HEL peptide was used to set the gate placement. Lymphocytes gates were selected based on forward and side scatter plots, dead cells were excluded using PI, and cells positive for APC anti-human CD8 antibody were selected as T cell gates from which GFP-positive T cells were assessed. **(B)** Flow cytometry gates show the percentage of GFP positive T cells before stimulation (*left*) and after stimulation (*right*) by co-culturing with APC and HEL peptide. Data from a representative experiment is shown. **APC:** Antigen presenting cell, **FSC:** Forward scatter, **GFP:** Green fluorescent protein, **HEL:** Hen egg lysozyme**, PI:** Propidium iodide, **SSC**: Side scatter.

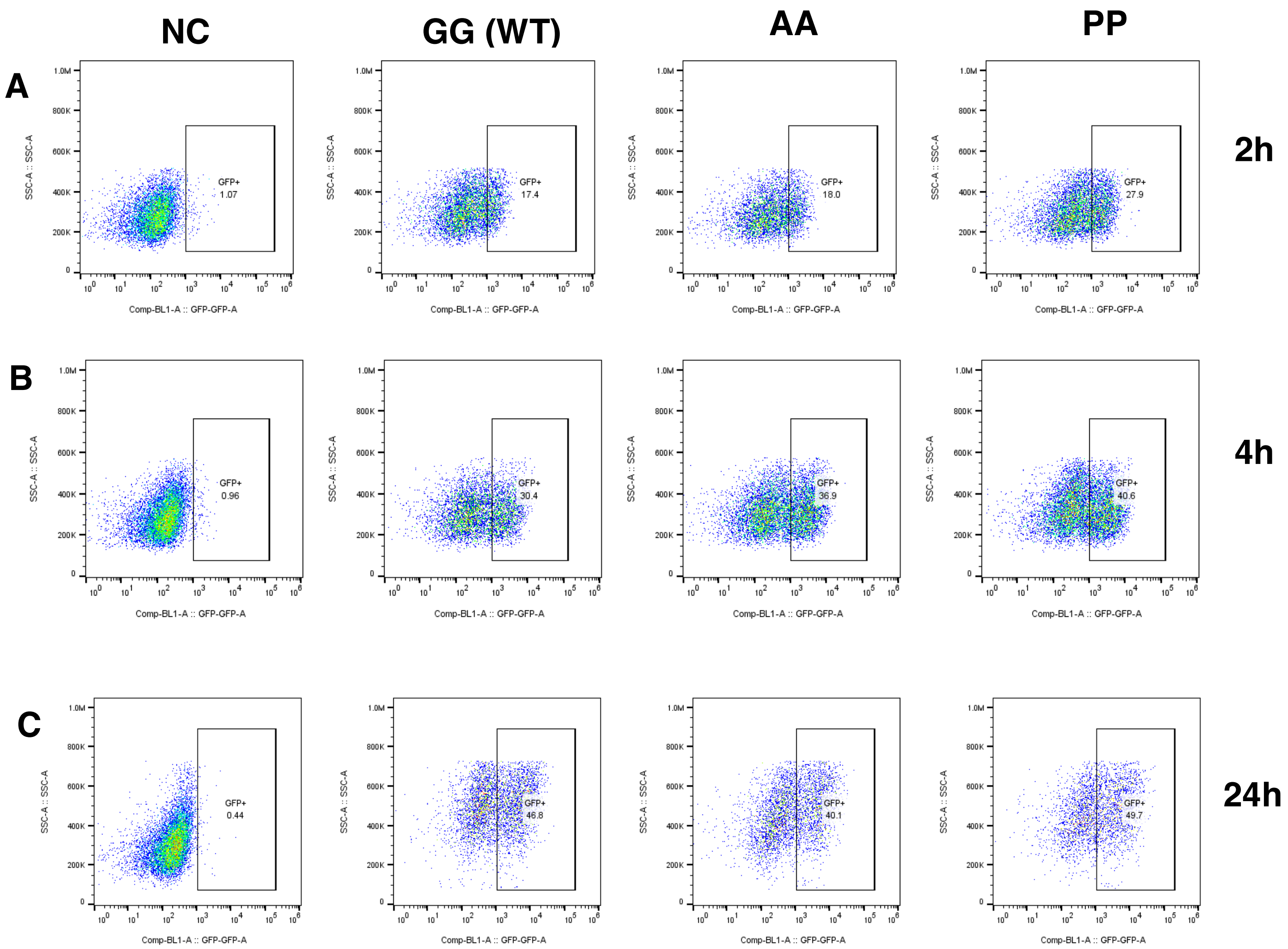


**Figure S8. Flow cytometer data analysis. (A-C)** Representative flow cytometry dot plots showing the percentage of GFP-positive T cells for the negative control, GG (WT), AA and PP cells after being stimulated with APC and 1 µg/ml HEL peptide for 2, 4, 24 h. **GG:** the wild-type 3A9 TCR with two glycine residues at positions 240 and 245 in the hinge region, **AA:** mutated hinge 3A9 TCR with alanines at positions 240 and 245, **PP:** mutated hinge 3A9 TCR with prolines at positions 240 and 245, **APC:** Antigen presenting cell, **GFP:** Green fluorescent protein, **HEL:** Hen egg lysozyme, **SSC**: Side scatter, **WT:** Wild type.

### Supplemental Bibliography

1. Mariuzza, R. A., Agnihotri, P. & Orban, J. The structural basis of T-cell receptor (TCR) activation: An enduring enigma. *Journal of Biological Chemistry* **295**, 914-925 (2020).
2. Natarajan, K. et al. An allosteric site in the T-cell receptor Cβ domain plays a critical signalling role. *Nature communications* **8**, 1-14 (2017).
3. Rangarajan, S. et al. Peptide-MHC (pMHC) binding to a human antiviral T cell receptor induces long-range allosteric communication between pMHC- and CD3-binding sites. *The Journal of biological chemistry* **293**, 15991-16005 (2018).
4. Kim, S. T. et al. The alphabeta T cell receptor is an anisotropic mechanosensor. *J Biol Chem* **284**, 31028-31037 (2009).
5. Das, D. K. et al. Force-dependent transition in the T-cell receptor β-subunit allosterically regulates peptide discrimination and pMHC bond lifetime. *Proc Natl Acad Sci U S A* **112**, 1517-1522 (2015).
6. Lanz, A. L. et al. Allosteric activation of T cell antigen receptor signaling by quaternary structure relaxation. *Cell Rep* **36**, 109375 (2021).
7. Newell, E. W. et al. Structural Basis of Specificity and Cross-Reactivity in T Cell Receptors Specific for Cytochrome c–I-Ek. *J Immunol* **186**, 5823-5832 (2011).
8. Dong, D. et al. Structural basis of assembly of the human T cell receptor–CD3 complex. *Nature* 1-21 (2019).
